# An information-borrowing Bayesian method for inferring high-dimensional population-level brain connectivity from small samples

**DOI:** 10.64898/2026.09.29.755278

**Authors:** Zhibin Pu, Chenhao Zeng, Shufei Ge

**Affiliations:** Institute of Mathematical Sciences, ShanghaiTech University, Shanghai, China

**Keywords:** brain connectivity analysis, high-dimensional covariance estimation, information borrowing, Hamiltonian Monte Carlo, ADHD

## Abstract

Estimating population-level dependence between brain regions is challenging when the number of regions greatly exceeds the number of individuals. We introduce an information-borrowing generalized Bayesian method for high-dimensional covariance estimation from region-level brain measurements. A divide-and-conquer parameterization separates marginal variances from correlation calibrations: an empirical *pilot* correlation matrix–by default, obtained by standardizing a sample covariance matrix (SCM)–provides a reference structure that is adaptively calibrated through region-specific parameters shared across pairwise correlations, reducing the number of free parameters from ***O*(*d***^***2***^) to ***O*(*d*)** (***d*** is the number of regions) while guaranteeing a positive-definite covariance through the inclusion of a ridge term. Horseshoe priors on the calibration parameters enable selective borrowing and shrinkage of pilot information. Under suitable conditions on pilot accuracy, we establish theoretical guarantees for covariance estimation while retaining scalability in ultra-high-dimensional settings via a low-rank likelihood representation and a Woodbury-accelerated Hamiltonian Monte Carlo algorithm. Simulations showed lower estimation error than the SCM, bagging and iteratively refined principal orthogonal complement thresholding (POETRY). Applications to a semi-synthetic cortical-thickness and ADHD resting-state fMRI data revealed significant group differences in the default-mode–visual network and somatomotor network–limbic network blocks after Benjamini–Hochberg correction.

## Main

The human brain is a complex system whose regions exhibit rich structural and statistical dependence, and estimating interregional dependency structures has become increasingly important in neuroimaging because such structures are associated with neuropsychiatric disorders [1]. Diffusion magnetic resonance imaging (dMRI) tractography provides a unique non-invasive tool for studying structural connectivity [2, 3] and structural-MRI (sMRI) studies have linked altered white matter integrity in frontal and temporal regions to psychiatric disorders like schizophrenia [4–6]. Beyond dMRI, modalities including functional magnetic resonance imaging (fMRI), magnetoencephalography (MEG), and electroencephalography (EEG) yield regional signals from which interregional statistical dependence–typically summarized as a network with regions of interest (ROIs) as nodes and dependence as edges–can be derived [7–12]. Such dependence networks are commonly analyzed through covariance or correlation matrix of regional measurements, which encodes pairwise population-level dependence[13–15], can be mapped to network edges, and serves as a generative blueprint for simulating realistic dependence patterns. Accurate covariance estimation is therefore foundational for network reconstruction and simulation[16, 17], yet remains statistically challenging in ultra-high-dimensional settings [18] where the number of regions *d* greatly exceeds the number of observations *n*; in this regime, the sample covariance matrix (SCM) is often singular and unreliable, making improved estimators essential.

Existing methods for large covariance matrix estimation fall into two primary strategies. One strategy leverages factor models by decomposing the covariance matrix into low-rank and sparse components, Σ = *L* + Φ, where covariance matrix Σ ∈ ℝ^*d*×*d*^, *L* is a low-rank component and Φ is a sparse matrix or a diagonal matrix. The principal orthogonal complement thresholding (POET) estimator is a mainstream example [19], extracting the low-rank latent factor component through principal component analysis and then thresholding the residual idiosyncratic covariance matrix. POET is optimization-free and computationally efficient, but its performance can deteriorate when the factor structure is weak or misspecified or the residual dependence is insufficiently sparse. The iteratively refined POET (POETRY) estimator improves POET by iteratively refining both the low-rank and the sparse components, reducing the bias of one-step separation procedure and achieving an improved convergence rate that attains the minimax lower bound for factor-based covariance estimation [20]. The other primary strategy uses variance–correlation decomposition, estimating the variance and correlation components separately and then integrating them to form an estimate of the covariance matrix, which is natural in a Bayesian framework. For example, [21] modeled standard deviations and correlations using inverse-Wishart and log-normal priors, but the parameter space expanded substantially and the inverse-Wishart prior led to slow convergence and high computational cost in high dimensions. [22] introduced more flexible priors on spheres with an adaptive Δ-spherical Hamiltonian Monte Carlo algorithm, but focused mainly on low-dimensional dynamic correlation processes. For high-dimensional covariance estimation, [23] proposed a bagging (bootstrap aggregating) calibration for the sample correlation matrix, which ensured positive definiteness with probability 1 in finite samples under mild conditions, yet only produces point estimates and cannot directly quantify uncertainty–a key limitation, since each sample correlation matrix may be singular and uncertainty quantification is often crucial for downstream interpretation.

Here, we propose an information-borrowing Bayesian (IBB) framework for ultra-high-dimensional covariance estimation in population-level interregional dependence analysis. IBB adopts a divide-and-conquer strategy for covariance modeling that separates marginal variance estimation from correlation modeling through information borrowing within a unified generalized Bayesian framework, guaranteeing a positive-definite covariance matrix. Central to this design is a pilot correlation matrix–supplied directly or obtained by standardizing an SCM or user-specified covariance matrix–that provides a template for interregional dependence, allowing IBB to borrow pairwise correlation information rather than estimate the dependence structure from scratch. IBB incorporates this reference structure through region-specific calibration parameters shared across pairwise correlations with horseshoe priors and models marginal variances jointly, while guaranteeing positive definiteness through the inclusion of a ridge term. This parsimonious construction reduces the number of free parameters for covariance estimation from *O* (*d*^2^) to *O* (*d*), allows distinct regularization of diagonal and off-diagonal components while retaining joint inference, and provides an interpretable framework for high-dimensional covariance estimation. We develop a scalable Hamiltonian Monte Carlo (HMC) algorithm that exploits the Woodbury matrix identity to reduce the computational cost of posterior sampling. Log-transforming the positive parameters ***σ***^2^, *τ*, and ***λ*** enables HMC sampling in an unconstrained parameter space. Across the simulation scenarios evaluated, IBB generally achieved lower estimation error than its competitors, including the SCM, bagging (BAG), and POETRY. Real-data analysis further demonstrates its practical utility for estimating population-level interregional dependence from brain imaging features.

## Results

### Overview of IBB

IBB estimates population-level interregional covariance and the corresponding correlations from ROI-level measurements in ultra-high-dimensional brain connectivity studies, where the number of regions can far exceed the number of individuals. In this setting, sample covariance and correlation matrices can be unstable or singular, compromising dependence estimation and downstream network inference. IBB adopts a divide-and-conquer covariance parameterization that separates marginal variance modeling from correlation calibration, with both components estimated jointly within a generalized Bayesian framework (as is shown in Fig. 1). IBB uses a pilot correlation matrix, supplied directly or obtained by standardizing the sample correlation matrix or a user-specified covariance matrix, as a template for interregional dependence. It jointly models marginal variances and calibrates pairwise correlations through region-specific parameters, while a ridge term ensures positive definiteness (Fig. 1**a**). A low-rank likelihood representation and Woodbury-accelerated HMC enable scalable posterior inference (Fig. 1**b**). Logarithmic transformations of the positive parameters ***σ***^2^, *τ*, and ***λ*** allow HMC to operate in an unconstrained space (Fig. 1**c**). A global–local horse-shoe prior on the calibration parameters adaptively shrinks weak or unstable pilot correlations while retaining stronger signals (Fig. 1**d**).

**Fig. 1:**
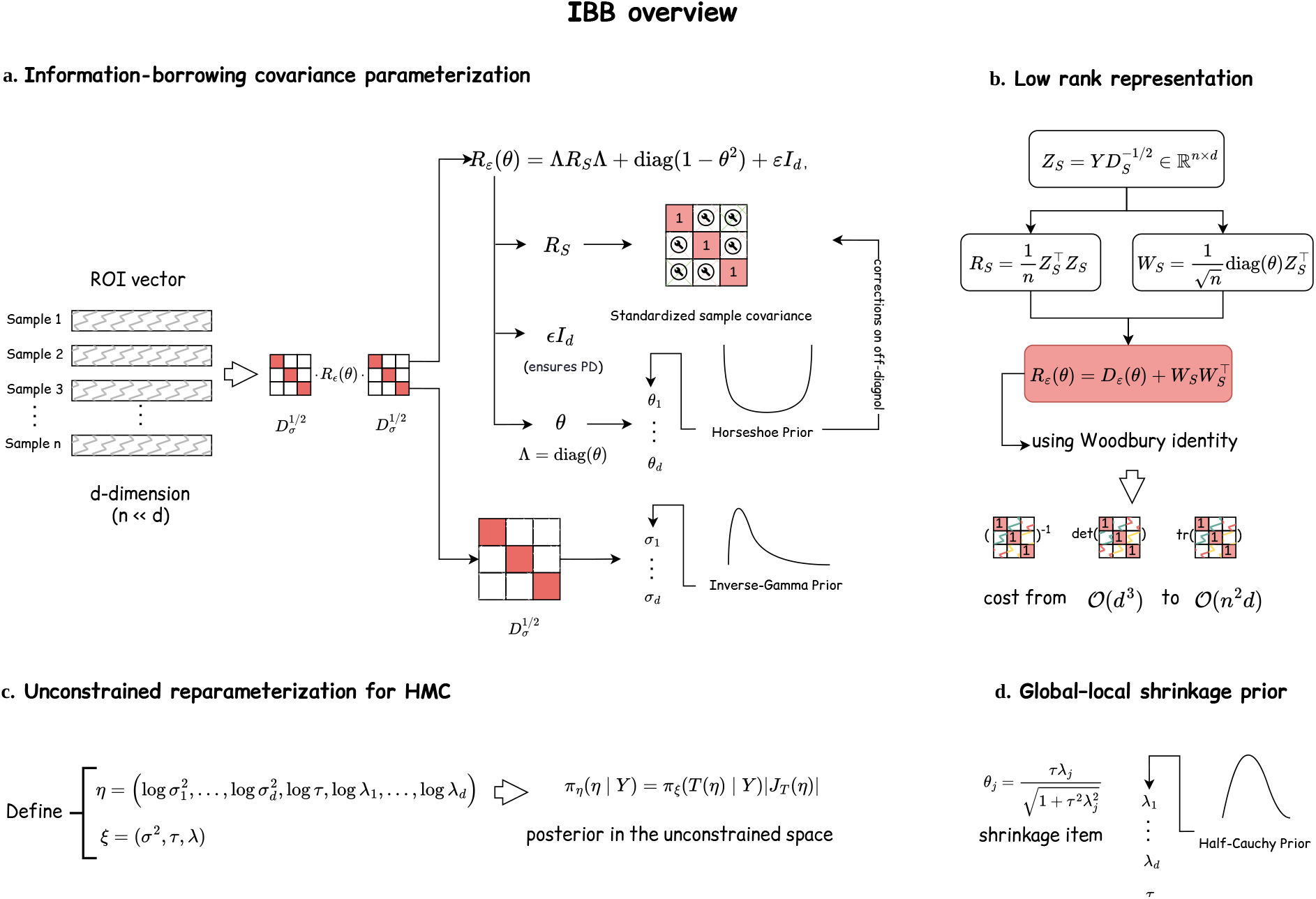
IBB method overview. IBB uses a pilot correlation matrix as a template for interregional dependence. **a**, IBB jointly models marginal variances and calibrates pairwise correlations through region-specific parameters, while a ridge term ensures positive definiteness. **b**, A low-rank representation of the likelihood, combined with the Woodbury identity, enables scalable posterior inference via HMC. **c**, Logarithmic transformations of the positive parameters ***σ***^2^, *τ*, and ***λ*** allow HMC to operate in an unconstrained parameter space. **d**, Global–local horseshoe priors on the calibration parameters adaptively shrink weak or unstable pilot correlations while preserving stronger signals.

The theoretical results establish structural and inferential guarantees for IBB and support efficient computation (Section Method). A ridge term with *ε >* 0 ensures that every induced correlation matrix is positive definite (PD) and, consequently, that the corresponding covariance matrix is also PD; see Theorem 1. Eigenvalue bounds established in Theorem 4 further provide conditioning guarantees that support numerically stable likelihood evaluation and posterior computation. Algebraic reformulations based on determinant and trace decompositions enable efficient evaluation of the log-determinants and quadratic forms required by the likelihood (Lemmas 2 and 3). Under the proposed prior specification, posterior propriety ensures that the posterior is well defined (Theorem 5). Complementing these foundational results, an accuracy guarantee is also established for the associated point estimator under active-node sparsity conditions (Theorem 6). Finally, Theorem 7 and Remark 1 theoretically justify the uniform enlargement of the off-diagonal entries of the pilot correlation matrix to correct systematic underestimation, showing that, under suitable conditions, the resulting cumulative perturbation either vanishes asymptotically or remains bounded.

Across all experiments, the IBB yields more accurate and stable connectivity estimates than conventional covariance methods and other competitive alternatives, yielding clearer group- and network-level effects.

### IBB improves covariance estimation in simulation studies

We generate *n* independent samples from a *d*-dimensional normal model *N* (0, Σ*), where Σ* has unit diagonal entries and a block, banded and Toeplitz covariance structure. We compared IBB with SCM, BAG, and POETRY in moderate- and ultra-high-dimensional settings by assessing recovery of the true covariance matrix Σ*. We measured overall estimation error using the Frobenius norm and the worst-direction error using the spectral norm. In the ultra-high-dimensional setting, we normalized the Frobenius error by the number of regions, 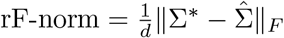. We also reported the proportion of estimated covariance matrices that were PD. To assess correlation calibration at the entry level, we compared the mean squared errors (MSEs) of the IBB and sample correlation estimates on two subsets: *P*, comprising entries for which the magnitude of calibration exceeded a specified threshold and *Q*, its complement. Performance on each subset was summarized using the relative MSE of IBB versus the sample correlation and the MSE reduction Δ = MSE^*S*^ − MSE^IBB^. A positive Δ indicates a lower MSE for IBB. Covariance constructions and evaluation metrics are detailed in the Supplementary Note 2.

In the moderate setting (*n* = 100, and *d* = 200–1,000), IBB achieved the lowest Frobenius- and spectral-norm errors across all covariance structures evaluated (Tables S1–S3 in the Supplementary; standard deviations are shown in parentheses). Among the competing methods, POETRY generally had lower errors than the BAG estimator under both metrics. This difference may reflect the shrinkage used by POETRY to balance the contributions of the SCM and the sparse residual, whereas BAG lacks an explicit shrinkage mechanism targeting sparsity. IBB, BAG and POETRY produced positive-definite covariance estimates in all evaluated moderate-dimensional settings.

In the ultra-high-dimensional setting (*d* = 15*n*), IBB generally achieved lower estimation errors than BAG and POETRY (Fig. 2**a**,**b**). SCM was omitted from these comparisons because its estimates were singular in this regime. Estimation errors are summarized as means across 100 simulation replications, with shaded bands indicating 95% confidence intervals for the mean.

**Fig. 2:**
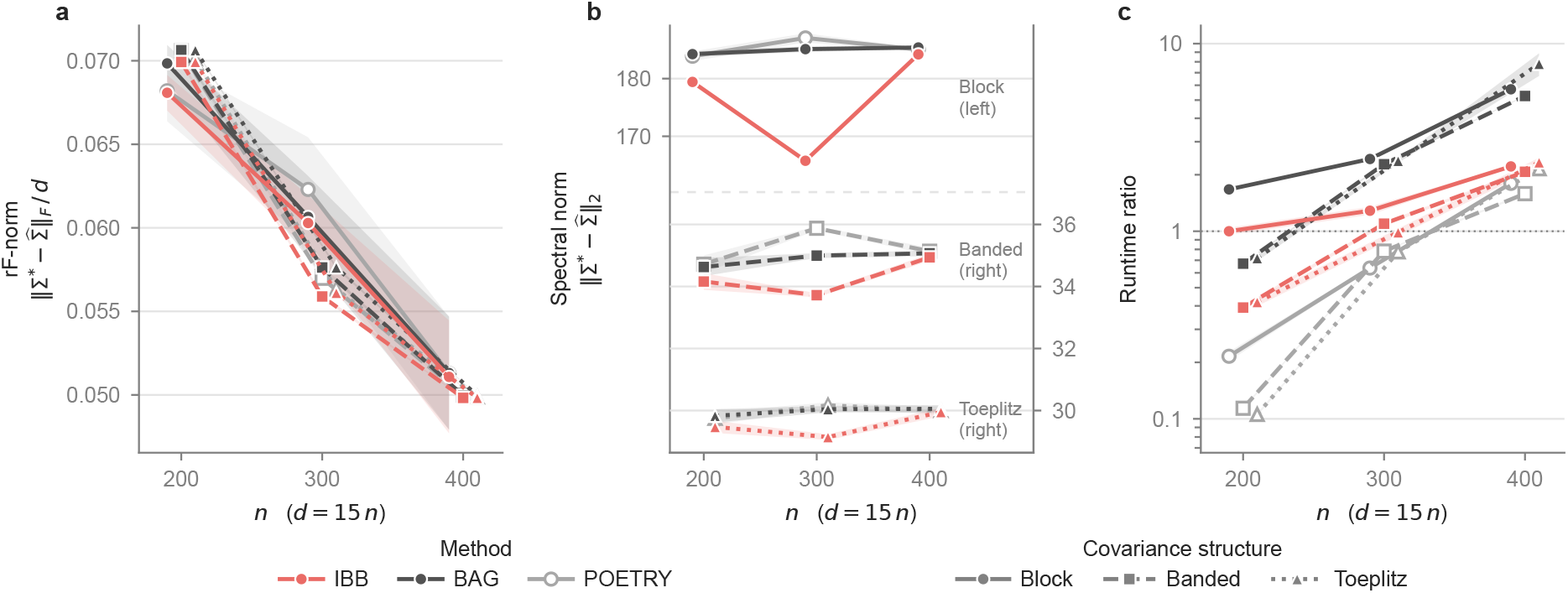
Estimation accuracy and computational efficiency in the ultra-high-dimensional setting. IBB, BAG, and POETRY are compared under block, banded, and Toeplitz covariance structures, with *n* ∈ {200, 300, 400} and *d* = 15*n*. **a**, Relative Frobenius-norm error (rF-norm), 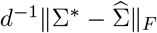, across sample sizes. Lines connect mean errors across 100 simulation replications, and shaded bands indicate 95% confidence intervals for the mean. **b**, Spectral-norm error, 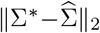, across sample sizes. Lines connect mean errors across 100 simulation replications, and shaded bands indicate 95% confidence intervals for the mean. The block structure is plotted against the left vertical axis, whereas the banded and Toeplitz structures are plotted against the right vertical axis. **c**, Relative runtime across sample sizes, normalized to the runtime of IBB under the block structure at *n* = 200. Runtime ratios are displayed on a logarithmic vertical axis, with a ratio of 1 corresponding to the baseline runtime. Colors distinguish covariance structures, and line styles distinguish methods throughout. Σ* and 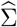 denote the true and estimated covariance matrices, respectively. Lower values indicate smaller estimation errors in **a**,**b** and lower computational cost in **c**.

For the rF-norm error (Fig. 2**a**), the three methods showed similar performance at *n* = 400 across the covariance structures, although IBB retained a small advantage. The rF-norm errors generally decreased as *n* and *d* increased together. This metric is the root mean squared error across the *d*^2^ covariance entries, so its behavior reflects both element-wise estimation accuracy and normalization by matrix size. For the block structure, strong correlations were concentrated within a few blocks, whereas most off-block entries were close to zero. Consequently, when off-block errors are small, differences in recovery within the correlated blocks may be less apparent in this entry-averaged metric.

The spectral-norm results provided a complementary assessment of worst-direction estimation error (Fig. 2**b**).The block results are read against the left vertical axis, whereas the banded and Toeplitz results are read against the right vertical axis. Errors for the banded structure remained relatively stable across sample sizes. For the block structure, errors varied less regularly with sample size, but IBB remained competitive and often achieved lower errors. Such non-monotonic patterns may reflect fluctuations in the dominant block-associated errors, because the spectral norm depends on the largest absolute eigenvalue of the symmetric error matrix. Together, the two error metrics indicated that IBB generally provided more accurate covariance estimates across the evaluated settings.

In terms of computational cost, IBB was faster than BAG in the ultra-high-dimensional settings evaluated, whereas POETRY had the lowest runtime (Fig. 2**c**). Runtimes were normalized to that of IBB under the block structure at *n* = 200 and are displayed on a logarithmic scale. IBB therefore offered a runtime advantage over BAG despite using sampling-based inference. POETRY was faster but had larger estimation errors than IBB in most settings, indicating a trade-off between computational cost and estimation accuracy.

Element-wise comparisons further showed that the MSE reduction relative to the sample correlations was larger on *P* than on *Q* across all covariance structures, dimensions and sample sizes evaluated (Tables S4 and S5 in the Supplementary). Thus, IBB achieved larger MSE reductions among entries receiving stronger calibration, suggesting that the model was particularly effective at correcting entries for which the empirical correlation requires stronger adjustment.

### Application to semi-synthetic cortical-thickness data

We conducted semi-synthetic cortical-thickness experiments based on sMRI-derived cortical-thickness measurements of 15 gifted children and 15 typically developing controls obtained from OpenNeuro (https://openneuro.org/datasets/ds002726/versions/1.0.1), each with 308 ROIs. We fitted probabilistic principal component analysis (PPCA) [24] separately to each group, fixing the latent dimension at 12, and used the resulting positive-definite PPCA covariance estimates as the reference matrices, Σ_*G*_ and Σ_*C*_ for the gifted and control groups, respectively. By combining the empirical group means with the reference matrices, we then generated semi-synthetic datasets with *n* = 50 samples per group under two data-generating models: (i) a Gaussian model and (ii) a multivariate *t*-distribution with *ν* = 10 degrees of freedom. IBB was compared with BAG and POETRY using the Frobenius and spectral-norm errors relative to the corresponding reference covariance matrix Σ* ∈ {Σ_*G*_, Σ_*C*_}, together with performance on a downstream Gaussian-likelihood-based classification task. Details are provided in Section Method.

IBB achieved lower Frobenius- and spectral-norm errors than BAG and POETRY under both the Gaussian and multivariate *t* models (Tables 1 and 2, respectively). These errors also had small empirical standard deviations, indicating more stable covariance recovery across replications. The ranking of the methods remained unchanged under the heavy-tailed *t* data-generating mechanism, suggesting that IBB’s covariance estimation advantage was retained under this departure from Gaussianity.

**Table 1:**
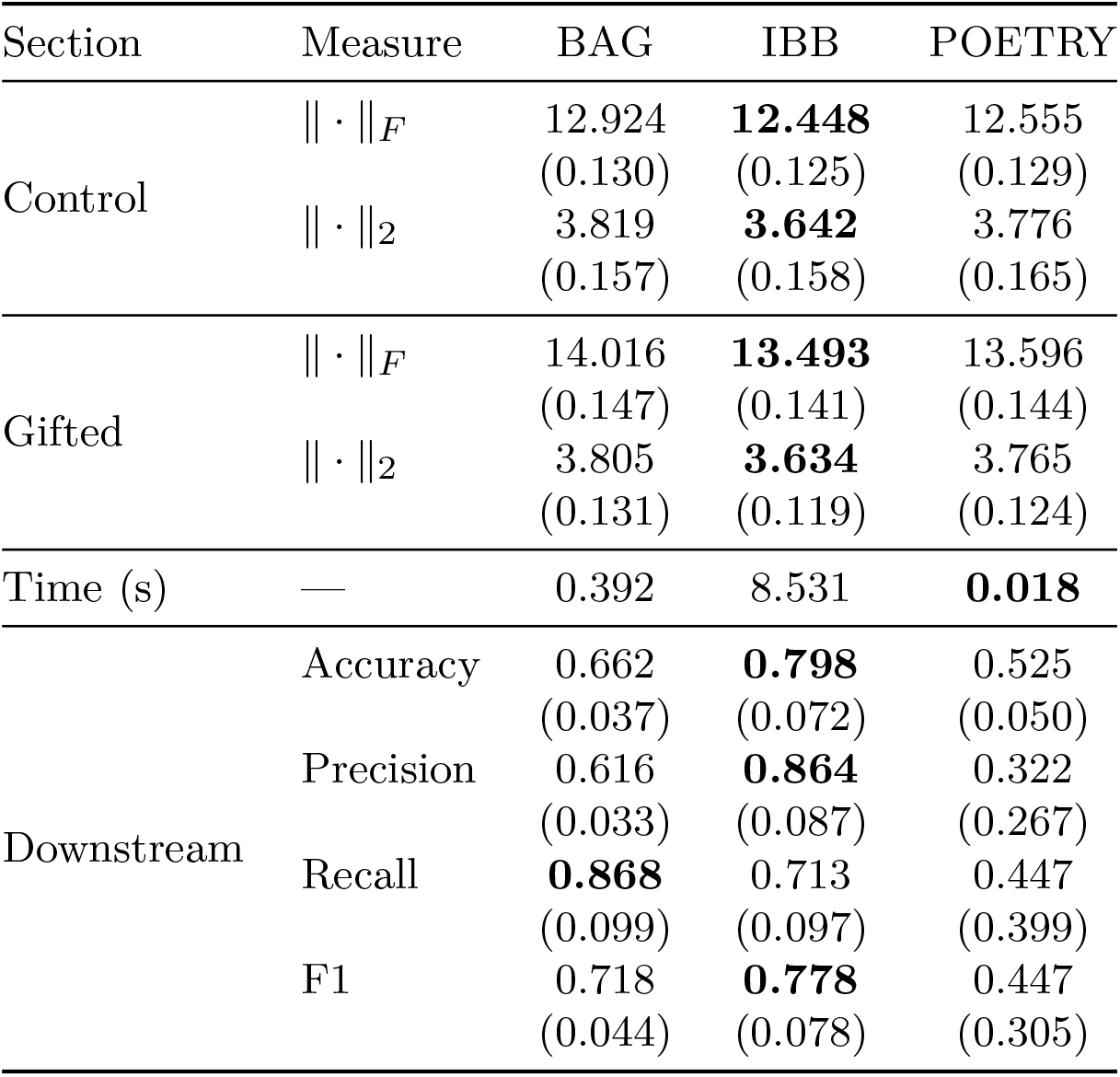
Averages and standard deviations (in parentheses) of the Frobenius norm and spectral norm for the gifted and control groups on the semi-synthetic Gaussian-distributed dataset, with *n* = 50, *d* = 308, and a train–test split of 2:1. The downstream analysis measures were calculated on the validation sets.

**Table 2:** Averages and standard deviations (in parentheses) of the Frobenius norm and spectral norm for the gifted and control groups on the semi-synthetic multivariate *t*-distributed dataset, with *n* = 50, *d* = 308, and a train–test split of 2:1. The downstream analysis measures were calculated on the validation sets.

| Section | Measure | BAG | IBB | POETRY |
| --- | --- | --- | --- | --- |
| Control | $\ \cdot\ _F$ | 18.300 | **17.911** | 18.309 |
|  |  | (1.643) | (1.792) | (1.856) |
| | $\ \cdot\ _2$ | 7.540 | **7.442** | 7.770 |
|  |  | (1.576) | (1.966) | (2.017) |
| Gifted | $\ \cdot\ _F$ | 19.855 | **19.433** | 19.863 |
|  |  | (1.788) | (1.959) | (2.029) |
| | $\ \cdot\ _2$ | 8.096 | **7.963** | 8.347 |
|  |  | (1.744) | (2.220) | (2.280) |
| Time (s) | — | 0.337 | 9.284 | **0.017** |
| Downstream | Accuracy | 0.626 | **0.704** | 0.504 |
|  |  | (0.053) | (0.077) | (0.085) |
|  | Precision | 0.616 | **0.751** | 0.331 |
|  |  | (0.060) | (0.108) | (0.279) |
|  | Recall | **0.755** | 0.628 | 0.360 |
|  |  | (0.194) | (0.078) | (0.406) |
|  | F1 | 0.658 | **0.680** | 0.493 |
|  |  | (0.081) | (0.078) | (0.311) |

In terms of computational cost, BAG and POETRY were substantially faster than IBB. For the Gaussian-likelihood-based downstream classification task, IBB achieved the highest accuracy, precision, and *F*_1_ score under both data-generating mechanisms (Tables 1–2). By contrast, BAG had the largest recall in both settings, indicating greater sensitivity (lower false-negative rate) at the expense of increased false positives. Overall, these results showed that IBB’s lower covariance reconstruction errors were accompanied by better performance on several classification metrics, at a higher computational cost. Details of the downstream classification are provided in Supplementary Note 3.

### Application to ADHD rs-fMRI reveals network-level differences ADHD-200 data and network annotation

We applied the proposed method to the publicly available ADHD-200 Sample, released through the International Neuroimaging Data-sharing Initiative (INDI) [25]. The ADHD-200 dataset aggregates resting-state fMRI (rs-fMRI), sMRI, and phenotypic information across multiple imaging sites. The primary analysis used data from the Peking University (PKU) cohort, and the New York University Child Study Center (NYU) cohort was subsequently used for validation. Our analysis focused on ROI-derived subject-level features and estimated group-level interregional dependence structures across individuals. Thus, the resulting correlations characterize covariation in ROI-derived subject-level features across subjects and should not be interpreted as conventional within-subject, time-series functional connectivity. After quality control, the PKU analysis included 78 participants with ADHD and 78 controls, with each participant represented by 955 ROI-derived features. Detailed information about the data and preprocessing procedures are provided in Section Method.

Because the target ROIs were not directly labeled according to anatomically named regions, we mapped them to the canonical networks defined by the **Yeo 7-network atlas** [26–28]. Specifically, the Yeo 7-network atlas partitions the cerebral cortex into seven canonical functional networks (CFNs) — the visual network (VIS), somatomotor network (SMN), dorsal attention network (DAN), ventral attention/salience network (VAN), limbic network (LIM), frontoparietal control network (FPN), and default mode network (DMN). These seven CFNs constitute a coarse yet neurobiologically meaningful representational framework for large-scale cortical functional organization. Broadly, VIS primarily governs visual perception and early sensory processing, whereas SMN is involved in sensorimotor integration and motor control. Top-down attention and goal-directed orienting to external stimuli are supported by the DAN, whereas salience detection and stimulus-driven attentional reorienting are handled by the VAN. Affective processing, memory, and visceral or autonomic functions are associated with the LIM, while cognitive control, executive regulation, and flexible decision-making are linked to the FPN. Internally directed cognition, self-referential processing, and memory-related processes are primarily managed by the DMN. The full names, abbreviations, and brief functional descriptions are summarized in the Supplementary Table S6.

### Network-level differences between ADHD and control groups

We next evaluated between-group differences at the network-block level. We first computed the ROI-level posterior mean correlation matrices for the control and ADHD groups, respectively. We then conducted block-level permutation tests for each pair of Yeo networks, testing the null hypothesis that inter-block dependency pattern did not differ between groups. To account for testing across multiple network pairs, we applied the Benjamini–Hochberg (BH) [29] to control the false discovery rate (FDR). Full details of the testing procedure are available in Section Method.

Figure 3 summarizes the network-level group comparison results, including the posterior mean correlation matrices for the control and ADHD groups and heatmaps of the block-wise permutation-based statistics expressed as −log(*p*) and BH-adjusted −log(*q*). Larger values indicate stronger statistical evidence against the null hypothesis. Note that the block statistics were computed from between-group correlation differences within each network pair. Therefore, the diagonal entries of the test heatmap could be different from each other. Under the natural-log transformation, the conventional significance threshold of 0.05 corresponds to −log(0.05) = 2.995. After FDR correction, significant between-group differences were identified for the DMN–VIS and SMN–LIM network pairs (*q <* 0.05).

**Fig. 3:**
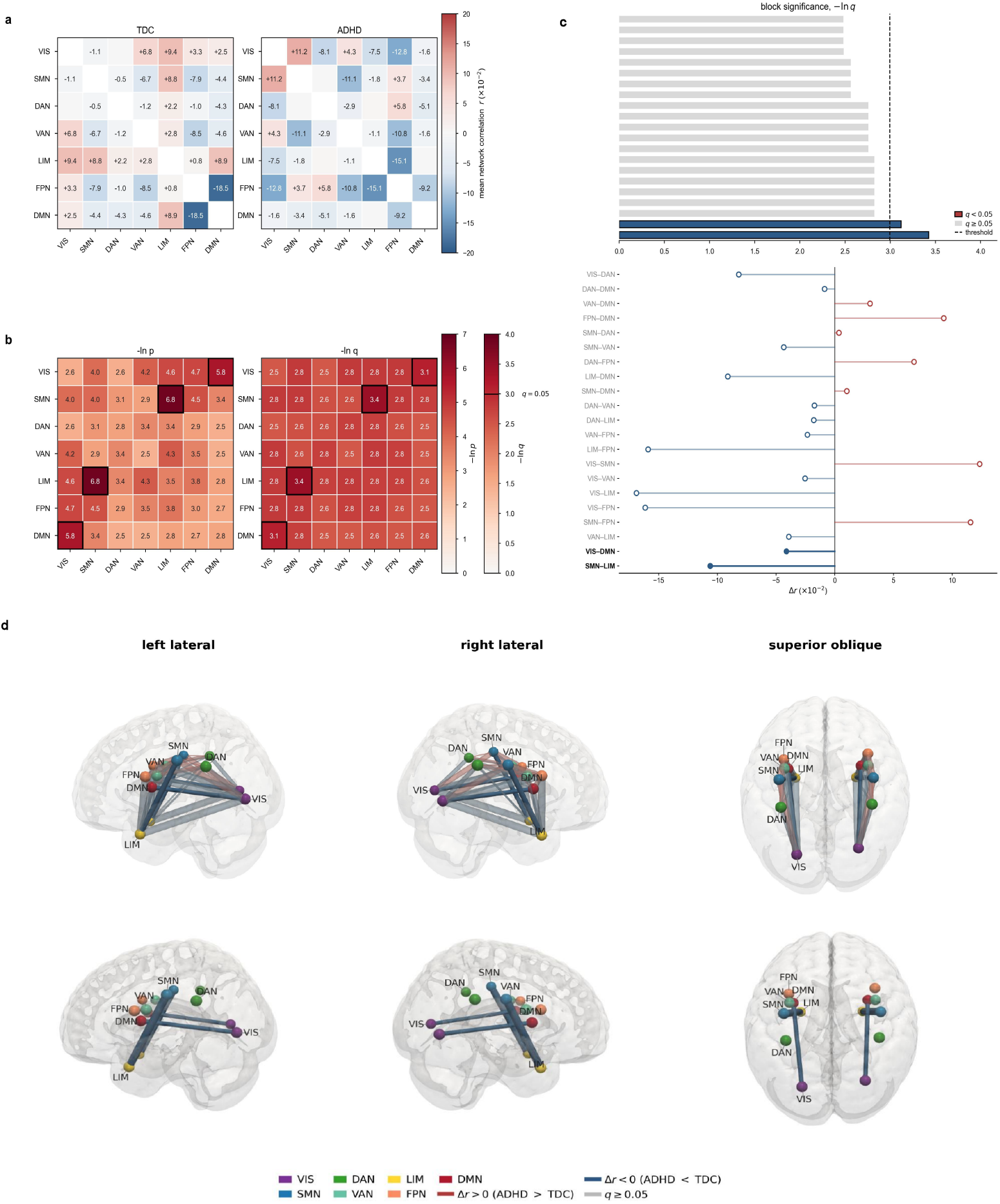
Network-level group comparison in the PKU ADHD-200 dataset. **a**, Block-wise posterior mean correlation matrix for the TDC group (left) and ADHD group (right). **b**, Heatmap of block-wise permutation test significance, shown as −log(*p*) (left) and heatmap of BH-FDR adjusted significance, shown as −log(*q*) (right). Network blocks correspond to pairs of Yeo 7 functional systems, including VIS, SMN, DAN, VAN, LIM, FPN, and DMN. In this panel, larger values indicate stronger evidence of group differences between the TDC and ADHD group. **c**, Block-level effect size. The upper bars give the BH-FDR adjusted significance of each network pair, − log(*q*); the lower lollipops give the signed difference in posterior mean correlation, Δ*r* = *r*_ADHD_ − *r*_TDC_. Color encodes the sign of Δ*r* (red positive, blue negative). The two FDR-significant blocks are drawn in bold with their pair names emphasized. **d**, Spatial rendering of ADHD-minus-TDC differences in network-level posterior mean correlations. The cortical surface was derived from the MNI152NL1i0n2009cAsym T1-weighted template after Gaussian smoothing (*σ* = 1 mm), using an isosurface approximating the grey matter– cerebrospinal fluid boundary, and shaded according to mean curvature. Nodes represent the centroids of the seven Yeo networks within each hemisphere (14 nodes in total). Tube color encodes the signed group difference, Δ*r*, and tube radius is proportional to |Δ*r* |. Each displayed network-pair effect is rendered once in each hemisphere for visualization only; the two renderings represent the same block-level comparison, not separate hemisphere-specific estimates or tests. The second row displays only network pairs with Benjamini–Hochberg-adjusted *q <* 0.05.

Although these results do not represent within-subject functional connectivity, the identified group-level dependence patterns are neurobiologically plausible in light of systems neuroscience findings. Interactions between the DMN and visual systems have been linked to the integration of perceptual input with internally generated semantic or mnemonic representations during reading, mind-wandering, and sustained emotional processing [30–33]. Similarly, the observed SMN–LIM difference is consistent with evidence that limbic and cingulate circuits interface affective valuation and interoceptive states with motor planning and bodily action, thereby supporting emotion–action integration [34, 35]. These correspondences provide contextual support for the detected network pairs, although they do not by themselves establish the underlying biological mechanisms.

Taken together, the results suggest that ADHD-related group differences in the analyzed ROI-derived features are distributed across interactions between large-scale networks rather than being confined to a single region. In particular, the detected DMN– VIS and SMN–LIM differences point to altered cross-network relationships involving perceptual–internally directed processing and affective–sensorimotor integration. These findings demonstrate the ability of the proposed method to translate high-dimensional regional dependence estimates into interpretable network-level comparisons while accounting for multiple testing.

### Validation in the NYU cohort

We further evaluated the proposed method using the NYU ADHD-200 cohort as an independent validation dataset, comprising 51 participants with ADHD and 51 controls, each represented by 955 ROI-derived features. The superior classification performance of IBB suggests that IBB not only provides a more accurate recovery of the case- and control-specific covariance structures, but also indicate that the covariance patterns estimated from the PKU data are reproducible in the NYU cohort and that the IBB captures these group-specific covariance differences more effectively than the BAG and POETRY. In addtion, after applying the same network-level analysis and multiple-testing corrections, the NYU results provided additional evidence for reproducible group-level patterns and broadly consistent network-level findings. The complete validation is reported in Supplementary Note 4, and the results are summarized in Supplementary Figure S1 and Supplementary Table S7.

## Discussion

Motivated by the estimation of population-level interregional dependence in brain imaging data, we proposed the IBB as a structured covariance-shrinkage framework for high-dimensional settings. The method addresses a setting in which the number of regions may be large relative to the available sample size, rendering direct estimation of the covariance structure unstable. Rather than regularizing each covariance entry independently, the IBB uses a parsimonious multiplicative representation to share information across entries through node-specific effects and a pilot covariance matrix. This formulation is intended primarily to recover stable, large-scale dependence patterns, rather than to identify or interpret individual edges in isolation. Meanwhile, a ridge term is incorporated into the covariance construction to guarantee positive definiteness, which is particularly important when the sample size is substantially smaller than the number of regions.

A central feature of the IBB is its use of the pilot covariance matrix as a data-informed structural reference. The method therefore does not estimate the high-dimensional dependence structure entirely from scratch, but instead retains and calibrates correlation patterns present in the pilot estimate. This information-borrowing mechanism is particularly useful in high-dimensional, low-sample-size scenarios, where the empirical covariance matrix may be unstable for direct use but may nevertheless contain valuable structural information about the overall organization of interregional dependence. The extent to which this strategy is effective depends on the pilot matrix capturing meaningful shared structure: if the pilot estimate is dominated by sampling variation or systematic bias, borrowing from it may also propagate those features into the posterior estimate. Although the ridge term ensures that the resulting covariance matrix remains PD, it does not by itself eliminate bias inherited from a misspecified or noisy pilot matrix.

The calibration and variance parameters provide a parsimonious mechanism for controlling this information borrowing. The region-specific calibration parameters are shared across pairwise correlations, determining how the dependence patterns represented in the pilot matrix are retained or modified, whereas the variance parameters account for differences in marginal variability across regions. Together with shrinkage-inducing priors, these components adapt the contribution of the pilot structure while discouraging unstable or overly complex covariance estimates. Compared with a fully edge-specific regularization, this structured calibration requires substantially fewer shrinkage parameters and may therefore offer greater numerical stability and a lower computational burden when the dimension is large.

To further improve computational scalability, the IBB employs a low-rank representation of the likelihood and a Woodbury-accelerated HMC algorithm. The Woodbury identity avoids repeated direct manipulation of the full high-dimensional covariance matrix during posterior computation, thereby making HMC more tractable in settings where the number of regions greatly exceeds the number of individuals. This computational strategy complements the *O* (*d*) parameterization: the latter reduces the number of calibration parameters, whereas the low-rank likelihood and Woodbury acceleration reduce the cost of posterior sampling.

Our simulation results showed that the IBB achieved lower covariance-estimation error than the SCM, BAG, and POETRY in the settings examined. These findings support the value of combining structured information borrowing with high-dimensional regularization, although they do not establish that the IBB will uniformly outperform these methods under all covariance structures or pilot specifications.

The pilot matrix, calibration parameters, variance parameters and ridge term serve complementary roles in the model. The pilot matrix provides an empirical template for the dependence structure, the calibration parameters regulate how strongly that template should be preserved or adjusted, the variance parameters ensure appropriate marginal scaling and the ridge term ensures that the resulting covariance estimate is PD. Their combination allows the IBB to stabilize covariance estimation while retaining an interpretable connection to the patterns observed in the data. This structure is well aligned with the applications in which inference concerns network-level organization or group-level differences in regional dependence. It may also provide a basis for incorporating group comparisons, covariate effects, hierarchical components, and network-level summaries. Such extensions would, however, require explicit methodological development and validation.

The parsimony of the IBB also defines its principal limitation. Because shrinkage is induced through node-specific effects, the regularization applied to a given covariance entry is determined jointly by its two incident nodes and is therefore not fully edge-specific. While this design improves parsimony and stability, it may be less effective when the underlying dependence structure contains highly localized edge-level irregularities that do not conform to broader node-level patterns. Therefore, IBB is a regularization method with structured constraints, rather than a fully flexible edge selection procedure. This limitation also constrains interpretation. A large posterior correction for a particular covariance entry should not be taken as evidence that the corresponding edge is uniquely important, because it reflects the joint contribution of the node-wise shrinkage terms. The method is therefore better suited to stabilizing and summarizing the dependence structure than to supporting definitive claims about isolated connections. Accordingly, the empirical findings are more appropriately viewed as reflecting differences in large-scale brain-feature dependence organization than as conclusive evidence for specific edge-level abnormalities. Any neurobiological interpretation should therefore remain cautious and primarily descriptive. An important direction for our future work is to augment the current node-wise formulation with an additional sparse edge-specific component.

In conclusion, the IBB provides a parsimonious approach to high-dimensional covariance estimation by combining a data-informed pilot structure with node-wise calibration and marginal variance modeling. The inclusion of a ridge term guarantees that the resulting covariance matrix is PD, while the low-rank likelihood representation and Woodbury-accelerated HMC algorithm support scalable posterior computation in ultra-high-dimensional settings. Its principal contribution is to stabilize the estimation of the large-scale dependence patterns without introducing a separate regularization parameter for every edge. The same structural constraint that enables this stability limits the method’s ability to represent isolated edge-level irregularities. Therefore, IBB is most appropriate when the scientific objective is to estimate and compare broad dependence structures, with future extensions needed for applications requiring more localized inference.

## Method

### Problem setup

We consider covariance estimation for population region-level measurements ***y***_*i*_ ∈ ℝ^*d*^, *i* = 1, …, *n*, in the high-dimensional regime *d* ≫ *n*. Our setting targets the underlying covariance matrix Σ ∈ *C*_*d*_, where *C*_*d*_ = {*A* ∈ ℝ^*d*×*d*^ : *A* ≻ 0} is the set of PD matrices. When *d* grows rapidly with *n*, direct use of the sample covariance is unstable and may fail to support reliable inverse-based inference; see [36] for related discussion on ultra-high-dimensional regimes.

### Information-borrowing covariance model

We propose IBB for (ultra) high-dimensional covariance estimation. Let *Y* = (*y*_*ij*_)_*n*×*d*_ denote the observed, centered data and let ***y***_*i*_ = (*y*_*i*1_, …, *y*_*id*_)^⊤^. We assume

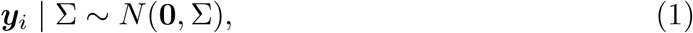

independently for *i* = 1, …, *n*.

### Information-borrowing covariance parameterization

The SCM,

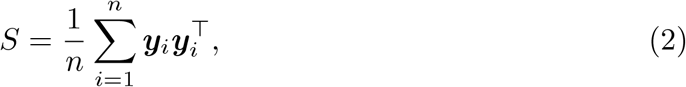

often reflects the global structure of Σ, yet can be singular in high dimensions and exhibits non-negligible errors both on the diagonal and off-diagonal entries. To leverage the global dependence pattern while correcting its unreliable parts, we embed the off-diagonal information from the normalized SCM into a variance–correlation decomposition.

Specifically, we parameterize Σ via

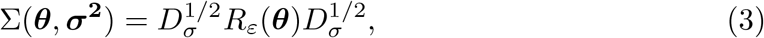

where

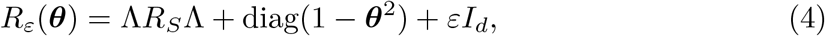

and

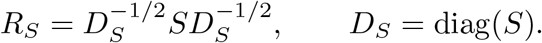

Here ***θ*** = (*θ*_1_, …, *θ*_*d*_)^⊤^ denotes node-wise calibration parameters with *θ*_*j*_ ∈ [0, 1] and Λ = diag(***θ***). Similarly, 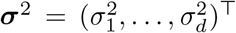 denotes variance parameters and *D*_*σ*_ = diag(***σ***^2^).

The ridge term *εI*_*d*_ is introduced to ensure that *R*_*ε*_(***θ***) is PD, and therefore Σ(***θ, σ***^2^) is PD. A detailed statement is given in Theorem 1 (Section Method). In addition, *ε >* 0 provides a uniform eigenvalue lower bound, particularly when *θ*_*j*_ → 1.

#### Relation to Bayesian separation strategy (BSS)

Equation (3) can be viewed as an adaptation of the BSS [21], where Σ = *D*^1*/*2^*RD*^1*/*2^ and the correlation matrix *R* and variance components are assigned separate priors. While fully Bayesian, such a formulation requires estimating *O* (*d*^2^) parameters for *R*, which can be computationally prohibitive and may raise identifiability concerns in the regime *n* ≪ *d*. In contrast, IBB replaces the full correlation parameterization by a calibrated, data-informed correlation structure through ***θ***, reducing the effective degrees of freedom to *O* (*d*) and enabling separate shrinkage for off-diagonal dependence while keeping variances strictly positive.

### Priors

To induce structured sparsity in the off-diagonal dependence, we impose a horseshoe-type global–local shrinkage prior on the calibration parameters. Specifically,

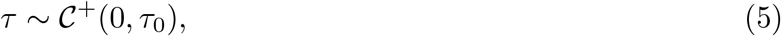

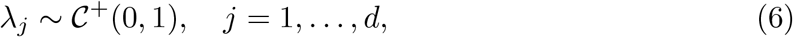

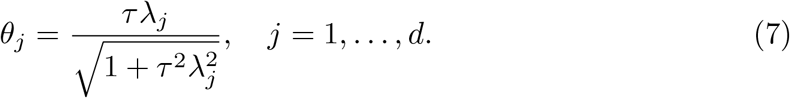

Here *C*^+^(0, ·) denotes a half-Cauchy distribution. Under this specification, ***θ*** is deterministically induced by (*τ*, ***λ***).

For the diagonal variance components, we treat ***σ***^2^ as unknown and assign independent inverse-gamma priors:

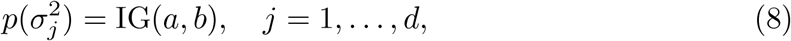

where IG(*a, b*) has shape parameter *a* and rate parameter *b*. This accounts for posterior uncertainty in variances and enables information sharing across ROI-derived features.

Since ***θ*** is determined by (*τ*, ***λ***) through Eq. (7), the model can equivalently be parameterized by (*τ*, ***λ, σ***^2^).

### Generalized posterior

Under the IBB framework specified above, the generalized Bayesian log-posterior for *τ*, ***λ***, and ***σ***^2^ takes the form

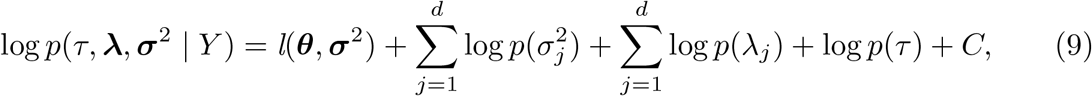

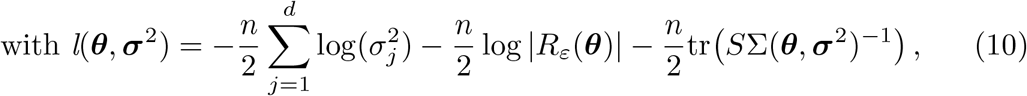

where *C* is a constant independent of the unknown parameters and tr(·) denotes the trace operator. The evaluation of the log-determinant and the quadratic/trace term is facilitated by the low-rank-plus-diagonal representation of *R*_*ε*_(***θ***); see Lemma 2 in Subsection Theoretical results of Section Method. In our studies and theory, we consider hyperparameters *τ*_0_ = 1, *a* = 3, *b* = 1, and *ε* = 10^−5^.

### Posterior computation via HMC

Closed-form posterior sampling is not available due to non-conjugacy between the priors and the Gaussian likelihood. We therefore use HMC, which augments the target posterior with momentum variables and generates proposals via Hamiltonian dynamics [37]. Compared with random-walk MCMC, HMC is typically more efficient in high-dimensional problems because it exploits gradient information to reduce random-walk behavior [38, 39].

#### Unconstrained reparameterization

To implement HMC on an unconstrained space, we apply logarithmic transformations to positive parameters ***σ***^2^, *τ*, and ***λ***. Define

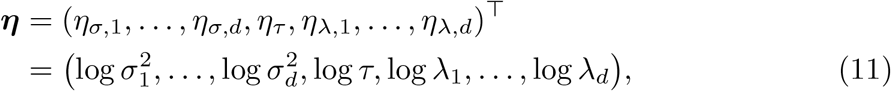

with inverse transformation

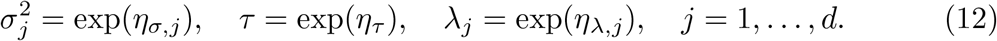

Let ***ξ*** = (***σ***^2^, *τ*, ***λ***) denote the constrained parameters and ***ξ*** = *T* (***η***) denote the inverse transformation. Under transformation, the posterior density in the unconstrained space is given by

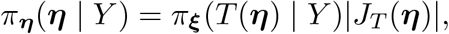

where *J*_*T*_ (***η***) is the Jacobian determinant. Since the mapping is component-wise exponential,

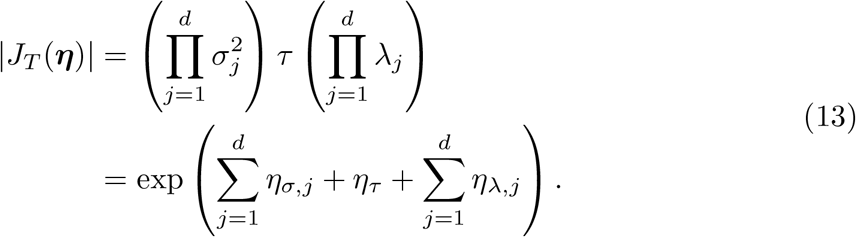

Accordingly, the log target density used in HMC equals

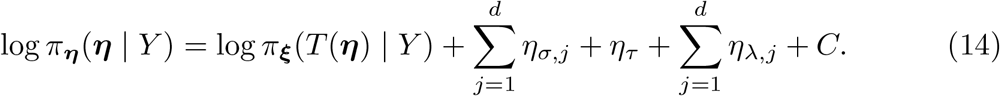

#### HMC algorithm

We use a leapfrog integrator with a Metropolis–Hastings correction to preserve the target distribution [38, 39]. The full procedure is summarized in Algorithm 1. Unless otherwise stated, we use *T* = 5000 iterations with 2500 warm-up iterations and 2500 posterior samples, and we adapt the step size to achieve an average acceptance probability close to 0.8.

### Computational efficiency

A direct evaluation of Eq. (10) requires computing determinants and inverses of Σ(***θ, σ***^2^), which has *O*(*d*^3^) complexity per HMC iteration when implemented naively. To enable scalable computation, we exploit the low-rank-plus-diagonal structure of *R*_*ε*_(***θ***) and use the Woodbury identity and the matrix determinant lemma [40].

Let 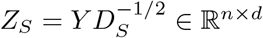 so that 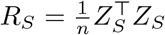. Then

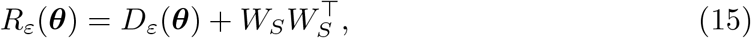

where

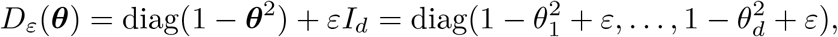

and

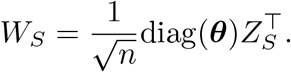

With this representation, the log-determinant and inverse terms can be evaluated using only *n* × *n* matrix operations, reducing per-iteration cost from *O* (*d*^3^) to *O* (*n*^2^*d*). The resulting expressions are

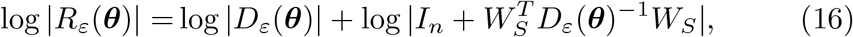

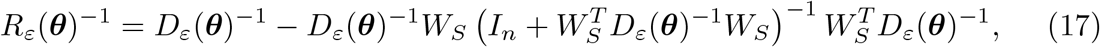

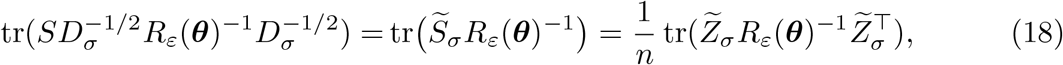

#### Algorithm 1

Hamiltonian Monte Carlo for the IBB framework

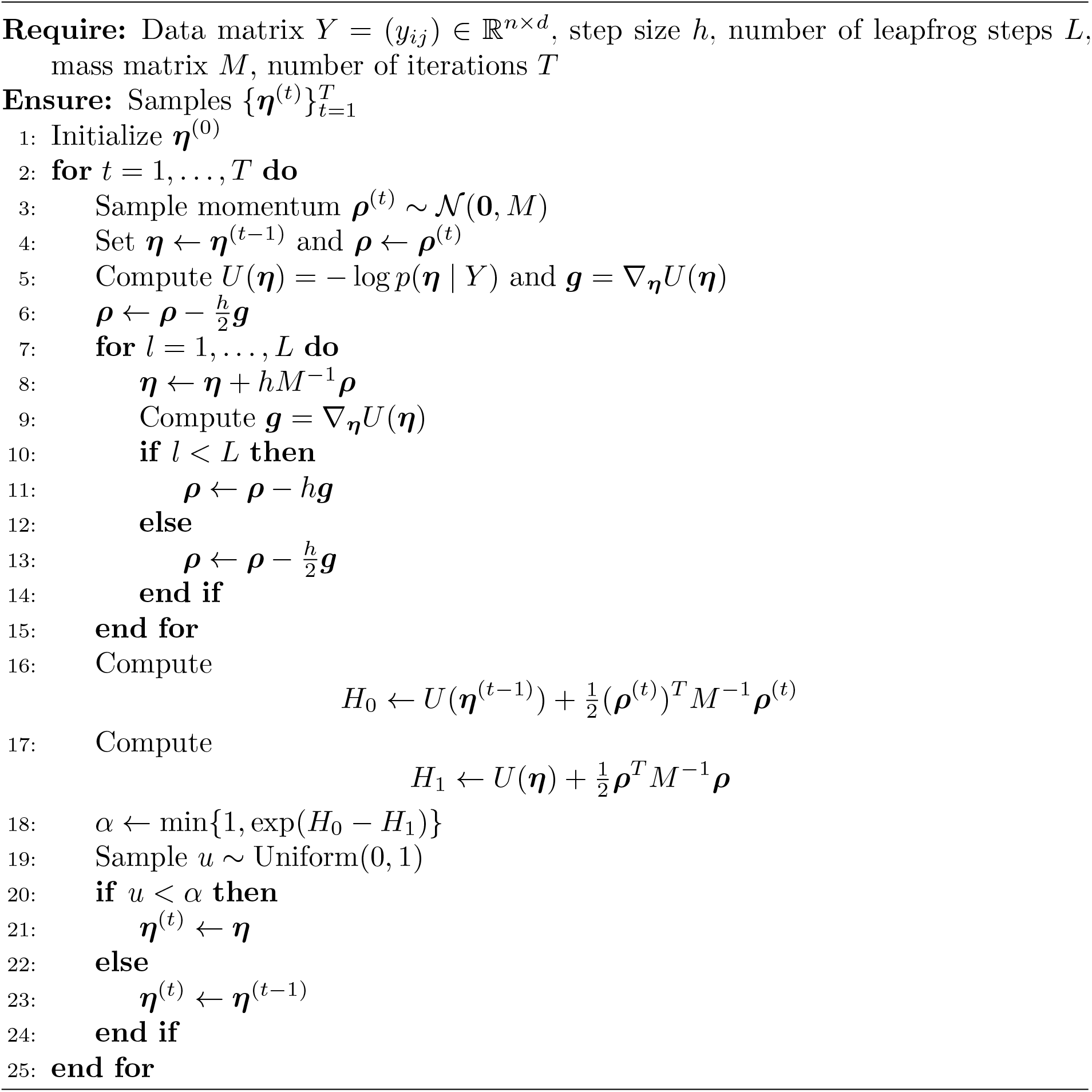

where 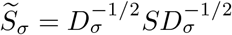 and 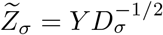. These computations are formalized in Lemma 2 and Lemma 3, and they also underpin several theoretical results.

### Theoretical results

We summarize the key theoretical properties of the proposed IBB framework. Specifically, these results establish that (i) the induced covariance family is well defined and PD (Theorem 1); (ii) eigenvalue bounds ensure numerical stability in likelihood evaluation (Theorem 4); and (iii) the determinant and trace terms admit scalable computational decompositions (Lemmas 2–3). We further establish that the induced posterior is proper under the proposed prior specification (Theorem 5) and that the associated point estimator satisfies an oracle-type approximation bound under active-node sparsity (Theorem 6). In addition, Theorem 7 and Remark 1 provide a theoretical justification for uniformly enlarging the off-diagonal entries of the pilot correlation matrix to correct systematic underestimation. Collectively, these results show that, under suitable conditions, the cumulative perturbation induced by this adjustment either vanishes asymptotically or remains bounded. For proofs and additional details, see Supplementary Note 1. The same statements remain valid if the pilot SCM *S* is replaced by a user-specified covariance (or correlation) matrix.

#### Theorem 1

*Let Y* = (*y*_*ij*_) ∈ ℝ^*n*×*d*^, *with rows* ***y***_*i*_ = (*y*_*i*1_, …, *y*_*id*_)^⊤^ ∈ ℝ^*d*^, *i* = 1, …, *n, and define*

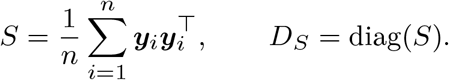

*Assume that the diagonal entries of S are strictly positive, so that* 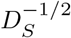 *is well defined. Let*

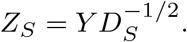

*For any* ***θ*** = (*θ*_1_, …, *θ*_*d*_)^⊤^ ∈ [0, 1]^*d*^, ***σ***^2^ = (*σ*^2^, …, *σ*^2^)^⊤^ ∈ (0, ∞)^*d*^, *and ε >* 0, *define*

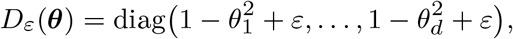

*and*

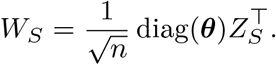

*Then the matrix*

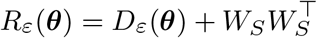

*is positive definite. Consequently, with*

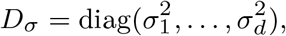

*the matrix*

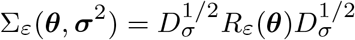

*is also positive definite*.

This result ensures that the IBB covariance estimator is well defined over the full parameter space.

#### Lemma 2

(Log-determinant decomposition) *Under the notation and assumptions of Theorem 1, the matrix R*_*ε*_(***θ***) *satisfies*

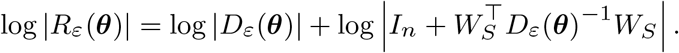

*Equivalently*,

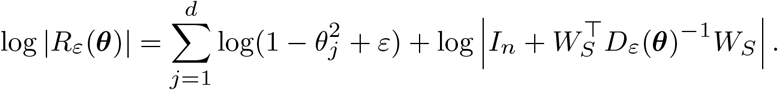

*Moreover, when n* ≪ *d, this decomposition reduces the computational cost of evaluating the log-determinant from the naive order O*(*d*^3^) *to*

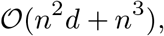

*which is dominated by O*(*n*^2^*d*) *whenever n*^3^ *is negligible relative to n*^2^*d*.

#### Lemma 3

(Trace decomposition) *Under the notation and assumptions of Theorem 1, the trace term satisfies*

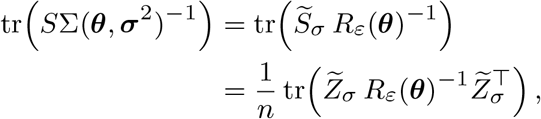

*where* 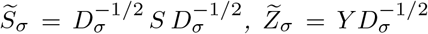. *Moreover, when n* ≪ *d, this decomposition reduces the computational cost of evaluating the trace term from the naive order O*(*d*^3^) *to*

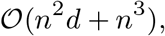

*which is dominated by O*(*n*^2^*d*) *whenever n*^3^ *is negligible relative to n*^2^*d*.

Lemmas 2, 3 indicate that the log-determinant and trace terms in Eq. (10) are computationally efficient in high-dimensional settings.

#### Theorem 4

(Conditioning bounds via Woodbury) *Under the notation and assumptions of Theorem 1, let* 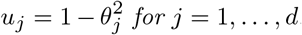. *For an arbitrary real square matrix A, let λ*_min_(*A*) *and λ*_max_(*A*) *denote its smallest and largest eigenvalues, respectively*.

*Then*

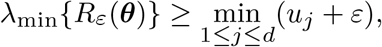

*and*

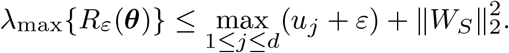

*Consequently, the condition number satisfies*

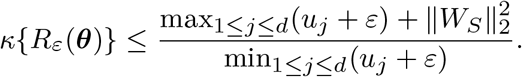

This bound provides a direct numerical stability guarantee for the likelihood evaluation and posterior computation.

#### Theorem 5

(Posterior propriety) *Under the notation and assumptions of Theorem 1, suppose that the probability model is specified as in Eq*. (1). *As described in Section Method, assume that the calibration parameters θ*_*j*_ *are determined by the global shrinkage parameter τ and the local shrinkage parameters λ*_*j*_, *j* = 1, …, *d, with the latter assigned proper half-Cauchy priors*.

*Further assume that the variance components satisfy*

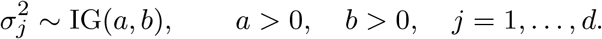

*Then, for any fixed ε >* 0, *the marginal normalizing constant*

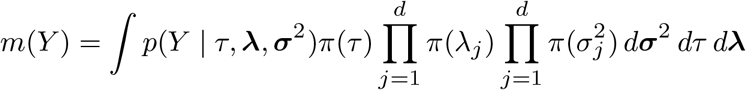

*is finite. Consequently, the posterior distribution is proper*.

This posterior propriety ensures that posterior-derived point estimators are well defined.

#### Theorem 6

(Oracle approximation bound under active-node sparsity) *Under the notation and assumptions of Theorem 1, let R*\* *denote the population correlation matrix and let R*_*S*_ *be its empirical counterpart (i*.*e*., *a borrowed pilot correlation matrix). Suppose that, on an event E*_*n*_, *the sample correlation matrix satisfies*

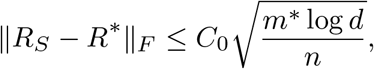

*where m*\* *denotes the number of nonzero off-diagonal entries in R*\*.

*Define the oracle approximation error*

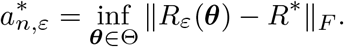

*Assume that* 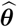 *satisfies the approximate projection condition*

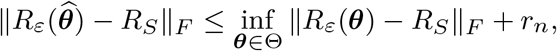

*where r*_*n*_ ≥ 0 *is the optimization or approximation tolerance. Then, on E*_*n*_,

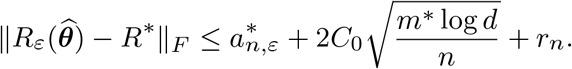

*In particular, if the support of R*\* *is induced by at most s active nodes, then*

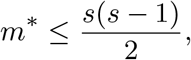

*and hence*

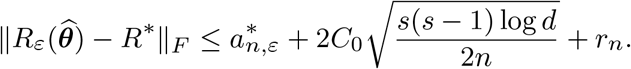

Theorem 6 should not be interpreted as a posterior contraction result. Rather, it provides a conditional approximation bound for a point estimator associated with the proposed data-dependent covariance family, under an accuracy event for the pilot correlation matrix *R*_*S*_.

The parameter vector ***θ*** determines the rescaling of the off-diagonal entries of the sample correlation matrix, with the corresponding rescaling factors ranging from 0 to 1. When the pilot matrix (e.g., the sample correlation matrix) systematically underestimates the population correlations, the calibrated parameters obtained without further adjustment may fail to recover the underlying population correlation matrix. To address this issue, we uniformly rescale, and in particular enlarge, the off-diagonal entries of the pilot matrix. Theorem 7 and Remark 1 show that this uniform rescaling controls the resulting cumulative perturbation: under suitable conditions, the additional perturbation either vanishes asymptotically or remains bounded.

#### Theorem 7

*Under the notation and assumptions of Theorem 6, define*

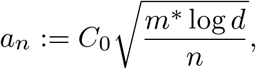

*where C*_0_ *>* 0 *is a constant. Let R*_*φ*_ *be a reconstructed correlation matrix associated with R*_*S*_ *(enlarged or shrunk) satisfying*

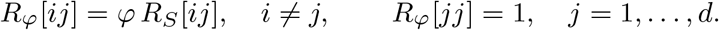

*Assume that the parameter φ >* 0 *satisfies*

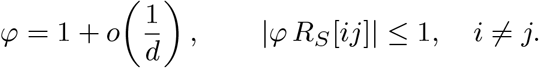

*Suppose that*

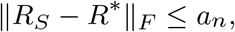

*and that the construction of R*_*φ*_ *yields the perturbation bound*

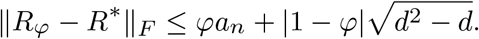

*Then, for all sufficiently large d, there exists a constant K >* 0 *such that*

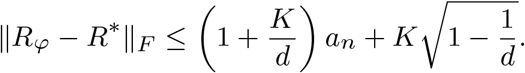

*More precisely, let*

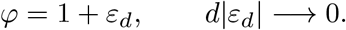

*Then*

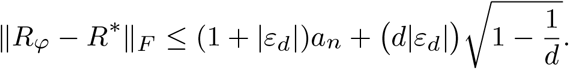

*Consequently*,

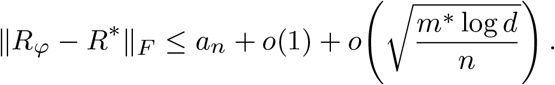

*In particular, if*

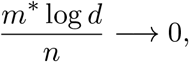

*then*

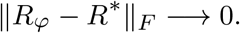

#### Remark 1

The Frobenius-norm error bound consists of two components:

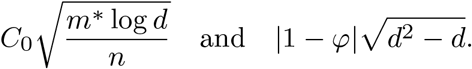

The first term represents the estimation error associated with the sample correlation matrix *R*_*S*_, where *n* denotes the sample size. The second term quantifies the perturbation induced by the rescaling parameter *φ* and depends on the matrix dimension *d*.

Since 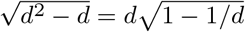 the perturbation term can be written as

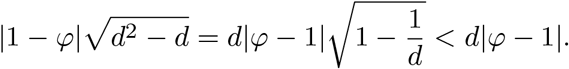

Thus, the condition *φ* = 1 + *o*(1*/d*) is sufficient to guarantee that *d*|*φ* − 1| → 0, so that the cumulative rescaling perturbation vanishes asymptotically. The parameter *φ* need not be greater than 1 or equal to 1; it may approach 1 from either side.

If this condition is not satisfied but the weaker condition 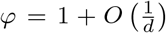 holds, then *d*|*φ* − 1| = *O*(1). In this case, the cumulative perturbation remains bounded.

### Simulation design

#### Simulation settings

Numerical experiments were conducted to compare the proposed IBB framework with the SCM and two representative competitors: BAG and POETRY.

BAG uses *d* resampling steps [23]. For POETRY, the number of factors *K* is selected automatically via an eigenvalue-ratio criterion [20]. For IBB, four MCMC chains with dispersed initial values were run, and convergence was assessed using potential scale reduction factors 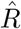 close to 1. All methods were implemented in Python 3.10 on a laptop with an Intel Core i7 (2.9 GHz) CPU.

For data generation, we sampled *n* independent observations from a *d*-dimensional multivariate normal distribution with mean zero and covariance matrix Σ*. The diagonal entries of Σ* were fixed at 1, and the off-diagonal entries were generated under three covariance structures—block, banded, and Toeplitz [41–43]. Details are provided in Supplementary Note 2.

Two regimes were considered: (i) a moderate-dimensional regime with *n* = 100 and *d* ∈ {200, 500, 1000}; and (ii) an ultra-high-dimensional regime where both *n* and *d* vary, with *n* ∈ {200, 300, 400} and *d* = 15*n*. When *d* is large, errors are reported using the relative Frobenius norm

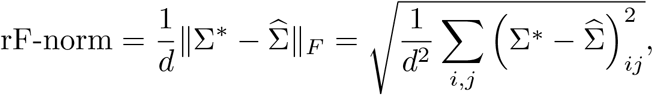

whereas the spectral norm is reported without normalization because it targets the worst-direction error.

#### Evaluation metrics

To evaluate an estimator 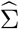, we use the following metrics: (i) Frobenius norm error 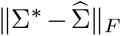; (ii) spectral norm error 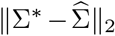; (iii) positive-definiteness rate, defined as the proportion of estimates 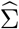 that are PD; (iv) MSE of the sample correlation matrix measured on specific index sets: 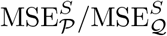; and (v) MSE of the IBB estimator measured on the same sets: 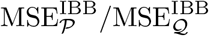. Here *P* denotes the index set of entries whose absolute differences exceed a threshold and *Q* is its complement; the above MSE definitions are given in Supplementary Note 2.2.

To summarize the relative improvement in MSE between the IBB estimator and the sample correlation, we further compute

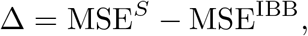

so that Δ *>* 0 indicates smaller MSE for IBB and Δ *<* 0 indicates smaller MSE for SCM. The above MSE reductions are reported based on the same *P* and *Q* defined in Supplementary Note 2.2.

### Semi-synthetic cortical-thickness experiment

#### Data description

In addition to the three specific covariance structures used in the simulation studies, we designed a semi-synthetic scenario using raw sMRI data from 15 gifted children and 15 typically developing controls—a dataset publicly available on the OpenNeuro platform (https://openneuro.org/datasets/ds002726/versions/1.0.1). For each subject, cortical thickness measurements were extracted from 308 brain ROIs, yielding a high-dimensional ROI-level representation with a small sample size (*n* = 15 per group). This setting is particularly challenging because it simultaneously faces the dual constraints of ultra-high dimensionality and limited data, which often results in ill-conditioned or singular sample covariance matrices, and because the true population-specific covariance structure is unknown, complicating direct assessment of covariance estimation accuracy [24].

#### Reference covariance construction

To obtain a positive-definite proxy ground truth while preserving realistic covariance structure, we constructed group-specific covariance matrices using PPCA. In particular, for each group, we fitted PPCA to the ROI-wise cortical thickness data and derived a positive-definite covariance matrix from the fitted PPCA model. The PPCA latent dimension was fixed at 12, which produced a well-conditioned reference covariance matrix. We denote the resulting reference covariances as the gifted-group covariance Σ_*G*_ and the control-group covariance Σ_*C*_. Group means were estimated empirically from the raw data and denoted by ***µ***_*G*_ and ***µ***_*C*_, respectively.

#### Data-generating mechanisms

Using the pairs (***µ***_*G*_, Σ_*G*_) and (***µ***_*C*_, Σ_*C*_), we generated semi-synthetic datasets with *n* = 50 samples per group under two distributional assumptions.

1. **Gaussian model**. For *i* = 1, …, *n*,

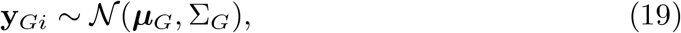

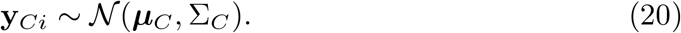
2. *t***-distribution model**. For *i* = 1, …, *n*,

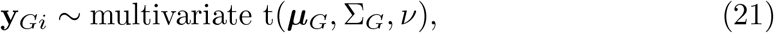

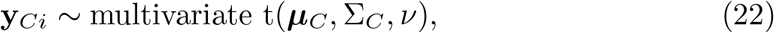

where *ν* denotes the degrees of freedom. We set *ν* = 10 to represent a moderately heavy-tailed alternative while keeping a well-defined second-moment structure.

#### Evaluation protocol

We compared IBB with BAG and POETRY using covariance reconstruction error measured by the Frobenius norm ∥ · ∥_*F*_ and the spectral norm ∥ · ∥_2_ between the estimated covariance 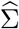 and the reference covariance Σ* ∈ {Σ_*G*_, Σ_*C*_}. For each data-generating mechanism, we repeated the data generation, estimation, and evaluation 50 times, and summarized performance using the mean and standard deviation.

To assess downstream utility, we additionally conducted a classification task based on a Gaussian likelihood. Classification metrics included accuracy, precision, recall, and *F*_1_ score; implementation details are provided in Supplementary Note 3.

### High-dimensional ADHD dataset

#### Data description

We applied the proposed model to the Peking University (PKU) ADHD-200 sample dataset to study group-level brain network structural differences between ADHD cases and typically developing controls (TDCs). The dataset was released by the International Neuroimaging Data-sharing Initiative (INDI) [25]. Because the original PKU sample exhibits substantial sex imbalance and prior studies suggest sex-related differences in functional brain organization and ADHD-associated neural patterns [44–46], we restricted the analysis to male participants. After randomly subsampling male controls to balance group sizes, the final analytical dataset contained 78 TDCs and 78 ADHD cases. For each participant, we extracted ROI-level features from 955 brain ROIs.

To further assess external generalizability, we repeated the same procedure on an independent NYU (New York University Child Study Center) ADHD-200 sample consisting of 51 male TDCs and 51 male ADHD cases, each represented by 955 ROI-level features [25].

#### Data preprocessing

The ADHD-200 dataset provides preprocessed resting-state fMRI regional time series. We summarized each participant by a *d*-dimensional feature vector obtained by averaging the time series within each ROI. To improve cross-subject comparability, each ROI feature was then z-score normalized. Rather than studying within-subject functional connectivity patterns, this ROI-averaged representation targets population-level dependence among brain-feature components and enables direct estimation of group-specific covariance/correlation structures.

#### Network-level differential analysis

We primarily examined group differences at the network-block level for the high-dimensional ADHA dataset. Specifically, each ROI is assigned to one of the seven CFNs via the mapping function *k* = *m*(*i*), where *i* = ∈ {1, …, 955}, and *k* 1, …, 7 indicates the index of the network to which it belongs. For each group *k*, we derived the posterior mean correlation matrix from the estimated covariance matrix using the separation-based representation in Eq. (3) in Section Method. Subsequently, the group-level difference matrix was defined as:

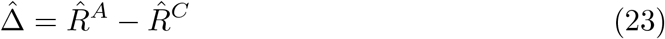

For any given network pair (*a, b*) (*a, b* ∈ {1, …, 7}), we define the set:

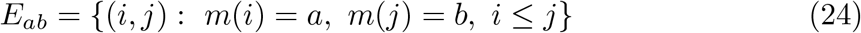

This set encompasses all ROI pairs belonging to the corresponding within- or between-network functional block. For each network pair (*a, b*), we summarized the group difference by the block statistic *T*_(*a,b*)_, defined as the average absolute difference:

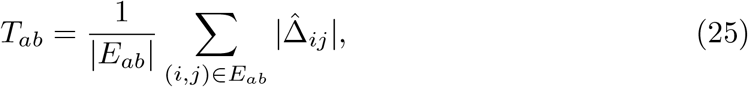

where |*E*_*ab*_| denotes the number of ROI pairs in the set *E*_*ab*_, and 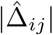 represents the (*i, j*)th entry of 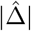. To assess the statistical significance, we performed a permutation test by randomly permuting group labels across subjects, refitting the model in each permuted dataset, and recomputing the block statistic 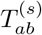, where *s* = 1, …, *S* denotes the permutation replicates (we set *S* = 1000). Specifically, we formally tested the null hypothesis *H*_0,*ab*_, which states that there was no statistically significant difference in the block-level inter-regional dependence patterns between the ADHD and control groups within network block (*a, b*)[47]. This hypothesis was quantified by the block-wise average absolute differences in their respective correlation matrices.

The corresponding permutation p-value [48, 49] was calculated as

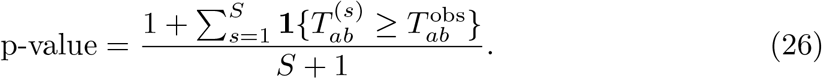

where 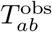 is the block-level observed statistic without permutation. Under *H*_0,*ab*_, the group labels are exchangeable, so permuting subject labels across the two groups provides a valid reference distribution for the block-level test statistic. The corresponding permutation p-value represents the probability, under the null hypothesis of no group difference, of observing a block-level statistic at least as large as that obtained from the original data. Thus, a relatively small p-value (e.g. *<* 0.05) indicates that the observed difference in block (*a, b*) is unlikely to have arisen solely by random grouping.

To account for multiple testing across all network pairs, we applied the BH to control the FDR and mitigate false-positive results [29]. Blocks with BH-adjusted p-values *q*_*ab*_ *<* 0.05 were considered significant, indicating a significant group difference in block (*a, b*).

## Acknowledgements

This project was supported by National Natural Science Foundation of China (12401383), and the startup fund of ShanghaiTech University. The authors also thank ShanghaiTech University for supporting this work through the startup fund and the HPC platform.

## Declarations

### Ethics approval

This study used publicly available, de-identified datasets from previously released repositories. No new human-participant data were collected by the authors.

### Consent to participate

Not applicable.

### Consent for publication

Not applicable.

### Data availability

The datasets analyzed in this study are publicly available from the repositories cited in the manuscript.

### Code availability

Code is available at https://github.com/ZhibinPU/IBB.git.

### Author contributions

Zhibin Pu: methodology, coding, formal analysis, writing— original draft, review and editing. Chenhao Zeng: data curation, investigation, formal analysis, writing—review and editing. Shufei Ge: conceptualization, supervision, methodology, writing—original draft, review and editing, funding acquisition.

## Supplementary Information

### Supplementary Figures

**Fig. S1.**
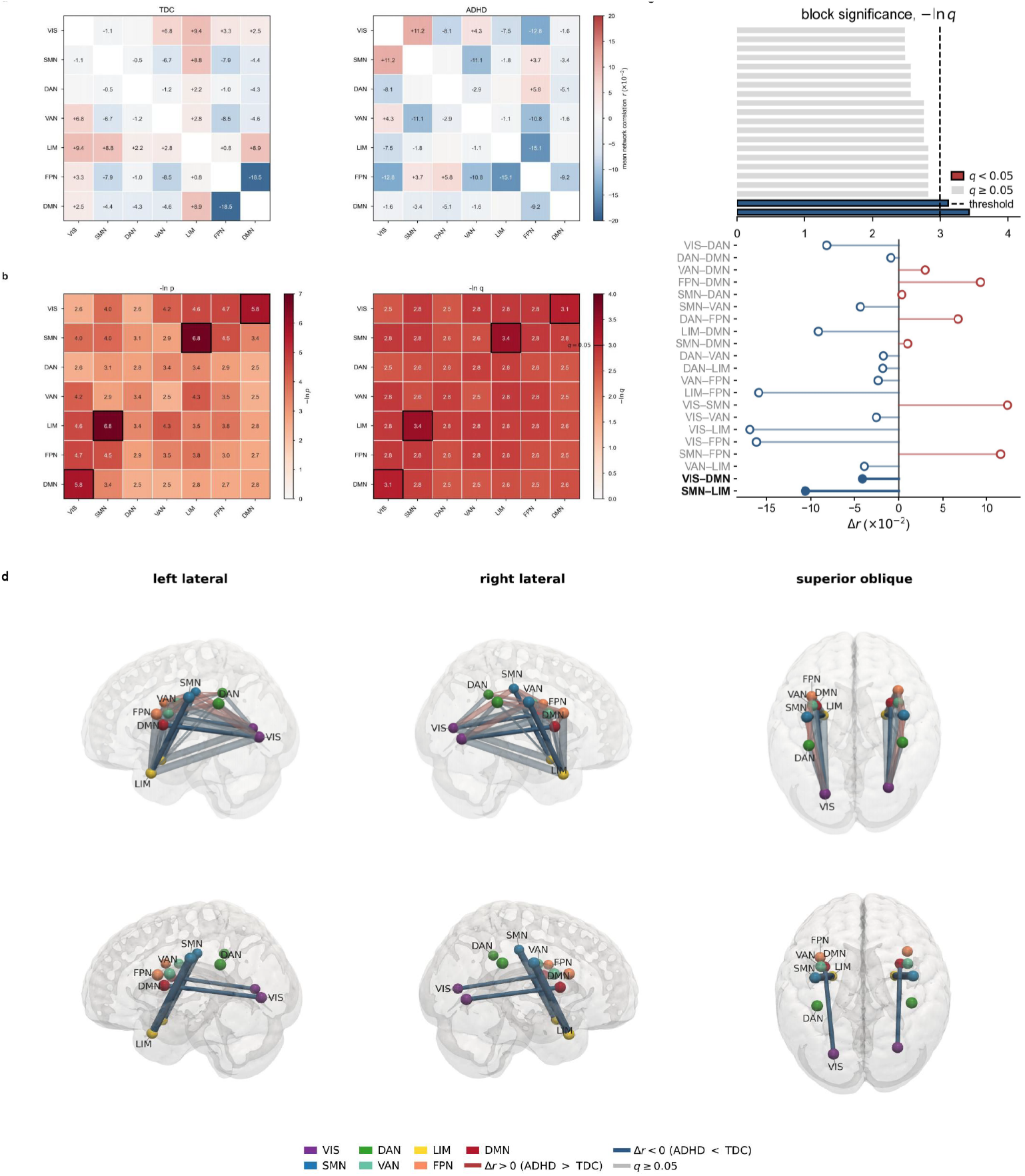
Network-level group comparison in the NYU ADHD200 dataset. **a**, Posterior mean correlation matrix for the TDC group (left) and ADHD group (right). **b**, Heatmap of block-wise permutation test significance, shown as −log(*p*) (left) and Heatmap of BH-FDR adjusted significance, shown as −log(*q*) (right). Network blocks correspond to pairs of Yeo 7 functional systems, including VIS, SMN, DAN, VAN, LIM, FPN, and DMN. In this panel, larger values indicate stronger evidence of group differences between the TDC and ADHD group. **c**, Block-level effect size. The upper bars give the BH-FDR adjusted significance of each network pair, −log(*q*); the lower lollipops give the signed difference in posterior mean correlation, Δ*r* = *r*_ADHD_ − *r* _TDC_. Color encodes the sign of Δ*r* (red positive, blue negative)3. The two FDR-significant blocks are drawn in bold with their pair names emphasised. **d**, Spatial rendering of ADHD-minus-TDC differences in network-level posterior mean correlations. The cortical surface was derived from the MNI152NLin2009cAsym T1-weighted template after Gaussian smoothing (*σ* = 1 mm), using an isosurface approximating the grey matter–cerebrospinal fluid boundary, and shaded according to mean curvature. Nodes represent the centroids of the seven Yeo networks within each hemisphere (14 nodes in total). Tube color encodes the signed group difference, Δ*r*, and tube radius is proportional to |Δ*r* |. Each displayed network-pair effect is rendered once in each hemisphere for visualization only; the two renderings represent the same block-level comparison, not separate hemisphere-specific estimates or tests. The second row displays only network pairs with Benjamini–Hochberg-adjusted *q <* 0.05.

### Supplementary Tables

**Table S1.**
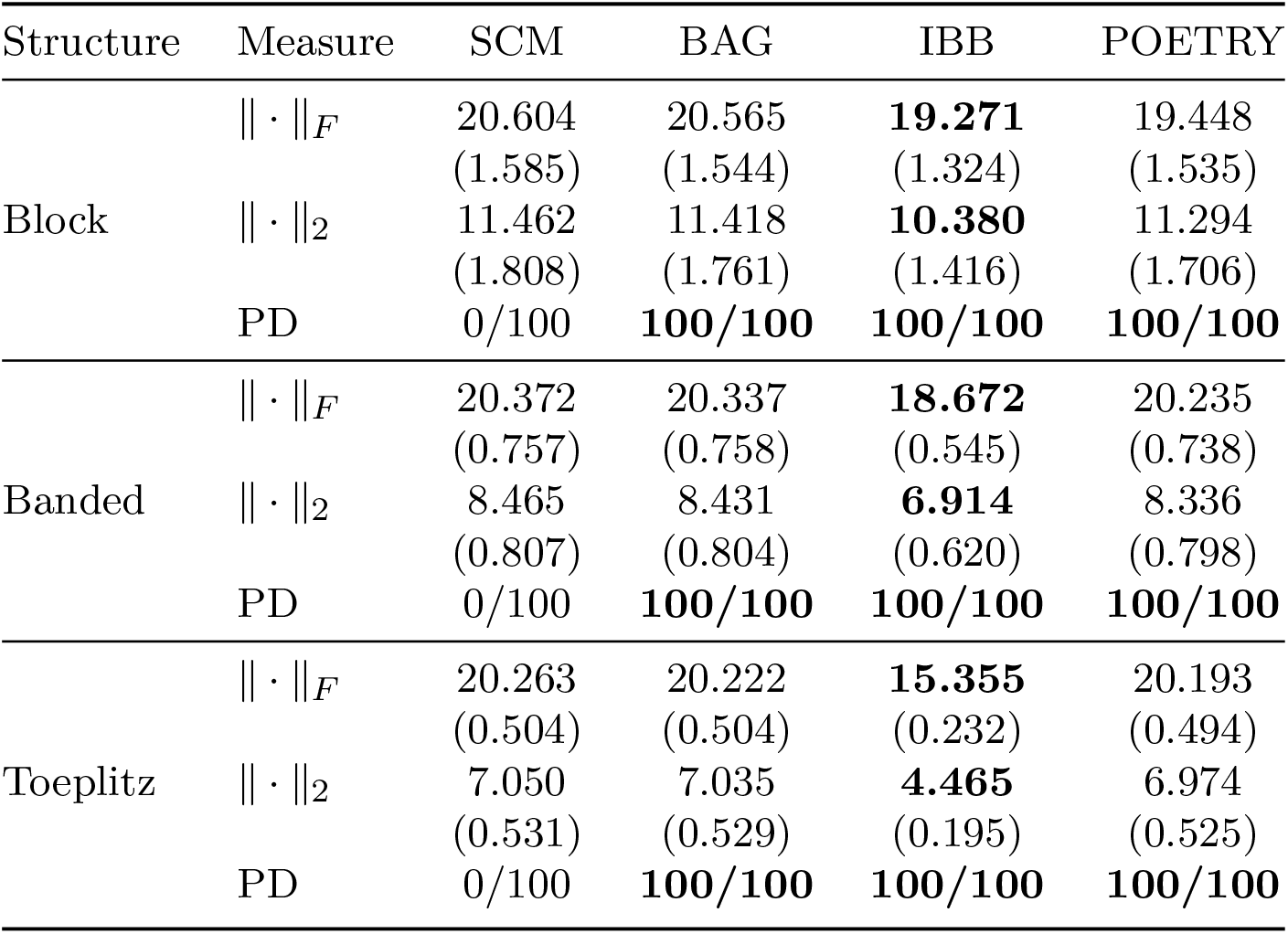
Mean estimation errors with standard deviations (in parentheses) over 100 replications when *n* = 100 and *d* = 200. Errors are reported in Frobenius norm and spectral norm.

| Structure | Measure | SCM | BAG | IBB | POETRY |
| --- | --- | --- | --- | --- | --- |
| Block | $\ \cdot\ _F$ | 20.604<br>(1.585) | 20.565<br>(1.544) | **19.271**<br>(1.324) | 19.448<br>(1.535) |
| | $\ \cdot\ _2$ | 11.462<br>(1.808) | 11.418<br>(1.761) | **10.380**<br>(1.416) | 11.294<br>(1.706) |
|  | PD | 0/100 | **100/100** | **100/100** | **100/100** |
| Banded | $\ \cdot\ _F$ | 20.372<br>(0.757) | 20.337<br>(0.758) | **18.672**<br>(0.545) | 20.235<br>(0.738) |
| | $\ \cdot\ _2$ | 8.465<br>(0.807) | 8.431<br>(0.804) | **6.914**<br>(0.620) | 8.336<br>(0.798) |
|  | PD | 0/100 | **100/100** | **100/100** | **100/100** |
| Toeplitz | $\ \cdot\ _F$ | 20.263<br>(0.504) | 20.222<br>(0.504) | **15.355**<br>(0.232) | 20.193<br>(0.494) |
| | $\ \cdot\ _2$ | 7.050<br>(0.531) | 7.035<br>(0.529) | **4.465**<br>(0.195) | 6.974<br>(0.525) |
|  | PD | 0/100 | **100/100** | **100/100** | **100/100** |

**Table S2.** Mean estimation errors with standard deviations (in parentheses) over 100 replications when *n* = 100 and *d* = 500. Errors are reported in Frobenius norm and spectral norm.

| Structure | Measure | SCM | BAG | IBB | POETRY |
| --- | --- | --- | --- | --- | --- |
| Block | $\ \cdot\ _F$ | 50.931<br>(4.042) | 50.806<br>(4.031) | **49.478**<br>(3.542) | 50.052<br>(3.934) |
| | $\ \cdot\ _2$ | 27.588<br>(5.115) | 27.476<br>(5.052) | **26.396**<br>(4.386) | 27.190<br>(4.933) |
|  | PD | 0/100 | **100/100** | **100/100** | **100/100** |
| Banded | $\ \cdot\ _F$ | 50.747<br>(1.167) | 50.555<br>(1.159) | **49.582**<br>(1.101) | 50.813<br>(1.149) |
| | $\ \cdot\ _2$ | 16.066<br>(1.162) | 15.995<br>(1.158) | **15.514**<br>(1.129) | 15.904<br>(1.150) |
|  | PD | 0/100 | **100/100** | **100/100** | **100/100** |
| Toeplitz | $\ \cdot\ _F$ | 50.529<br>(0.719) | 50.333<br>(0.715) | **41.065**<br>(0.592) | 50.104<br>(0.718) |
| | $\ \cdot\ _2$ | 13.483<br>(0.791) | 13.422<br>(0.781) | **10.077**<br>(0.633) | 13.448<br>(0.781) |
|  | PD | 0/100 | **100/100** | **100/100** | **100/100** |

**Table S3.**
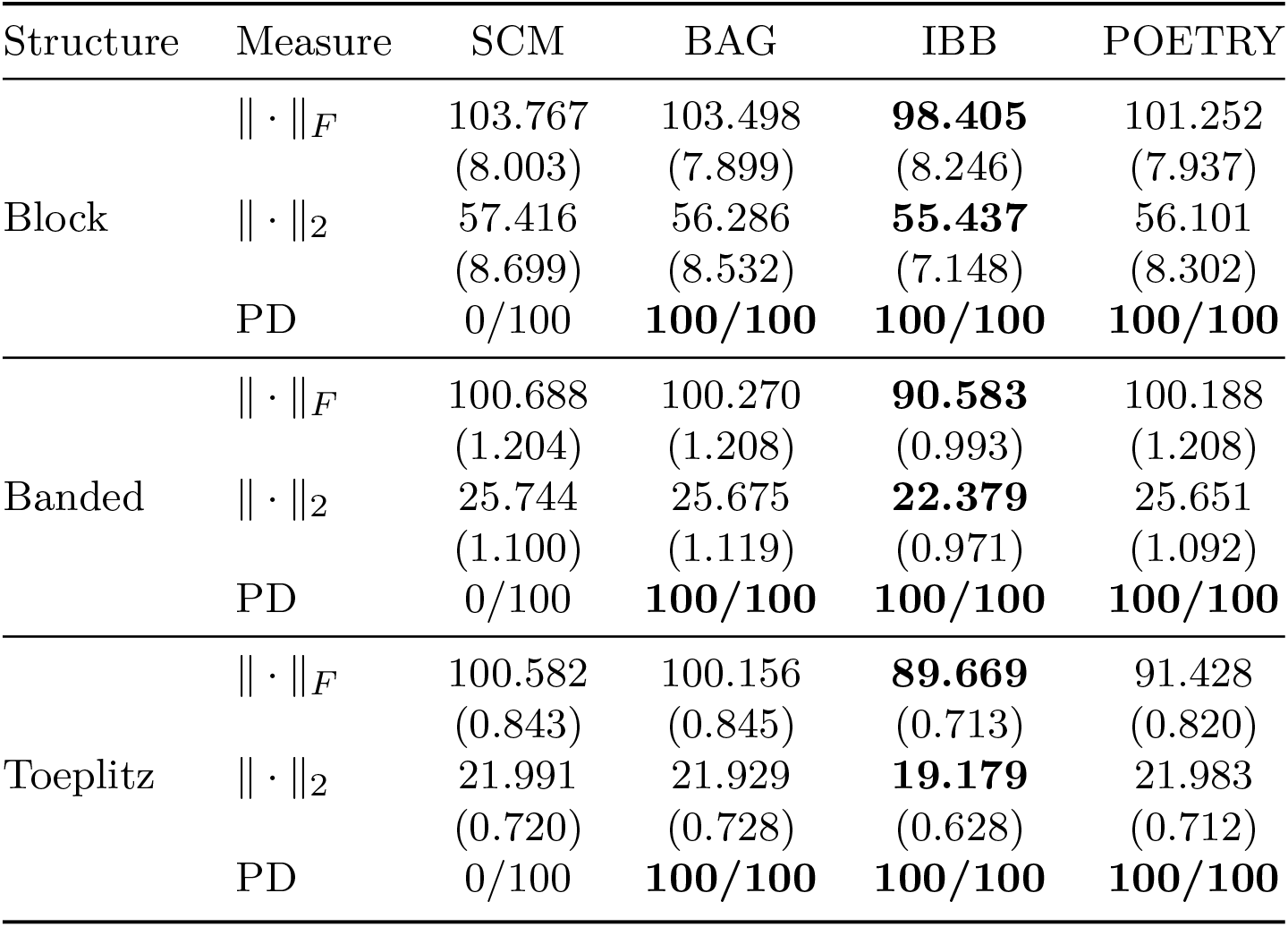
Mean estimation errors with standard deviations (in parentheses) over 100 replications when *n* = 100 and *d* = 1 × 10^3^. Errors are reported in Frobenius norm and spectral norm.

| Structure | Measure | SCM | BAG | IBB | POETRY |
| --- | --- | --- | --- | --- | --- |
| Block | $\ \cdot\ _F$ | 103.767<br>(8.003) | 103.498<br>(7.899) | **98.405**<br>(8.246) | 101.252<br>(7.937) |
| | $\ \cdot\ _2$ | 57.416<br>(8.699) | 56.286<br>(8.532) | **55.437**<br>(7.148) | 56.101<br>(8.302) |
|  | PD | 0/100 | **100/100** | **100/100** | **100/100** |
| Banded | $\ \cdot\ _F$ | 100.688<br>(1.204) | 100.270<br>(1.208) | **90.583**<br>(0.993) | 100.188<br>(1.208) |
| | $\ \cdot\ _2$ | 25.744<br>(1.100) | 25.675<br>(1.119) | **22.379**<br>(0.971) | 25.651<br>(1.092) |
|  | PD | 0/100 | **100/100** | **100/100** | **100/100** |
| Toeplitz | $\ \cdot\ _F$ | 100.582<br>(0.843) | 100.156<br>(0.845) | **89.669**<br>(0.713) | 91.428<br>(0.820) |
| | $\ \cdot\ _2$ | 21.991<br>(0.720) | 21.929<br>(0.728) | **19.179**<br>(0.628) | 21.983<br>(0.712) |
|  | PD | 0/100 | **100/100** | **100/100** | **100/100** |

**Table S4.**
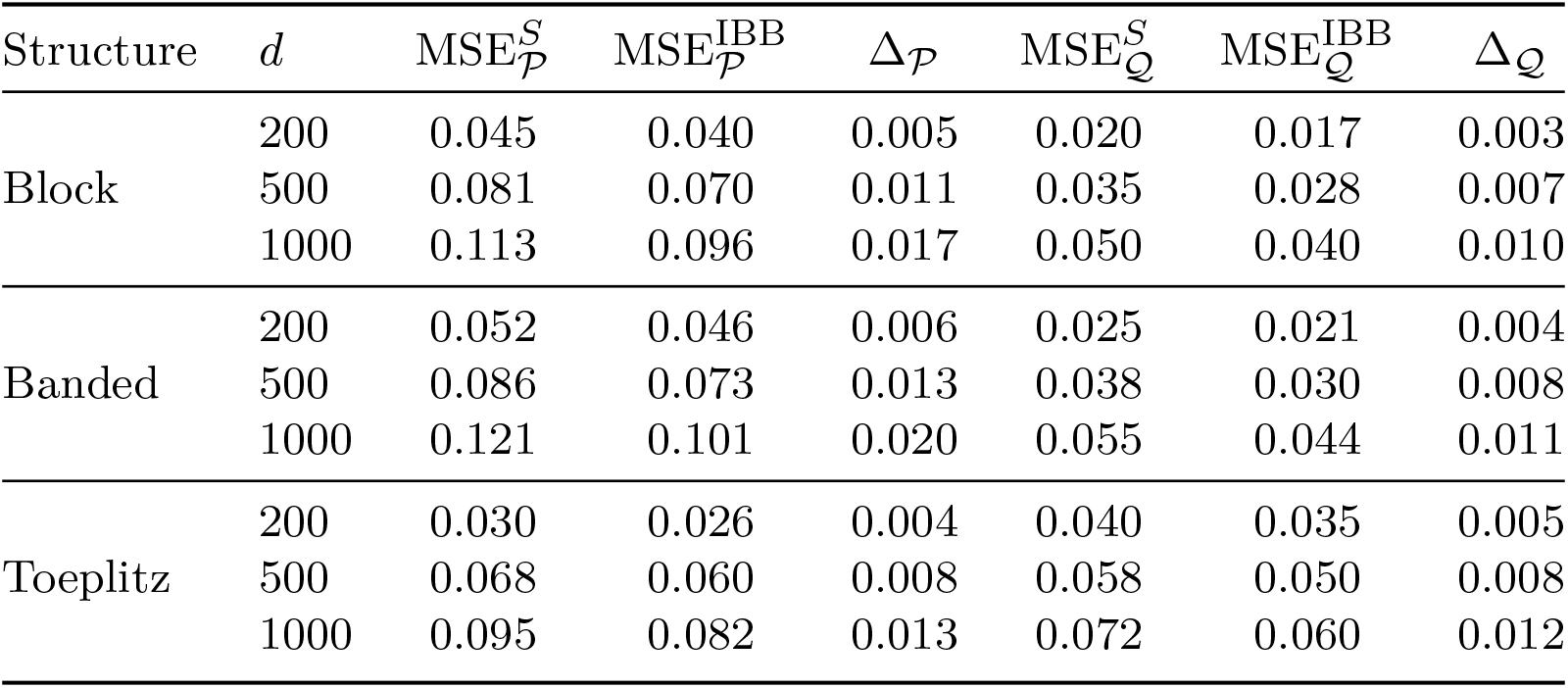
Element-wise mean squared errors on the correction-relevant subset *P* and its complement *Q* under the moderate-dimensional design with fixed sample size *n* = 100 and dimensions *d* ∈ {200, 500, 1 × 10^3^}.

**Table S5.**
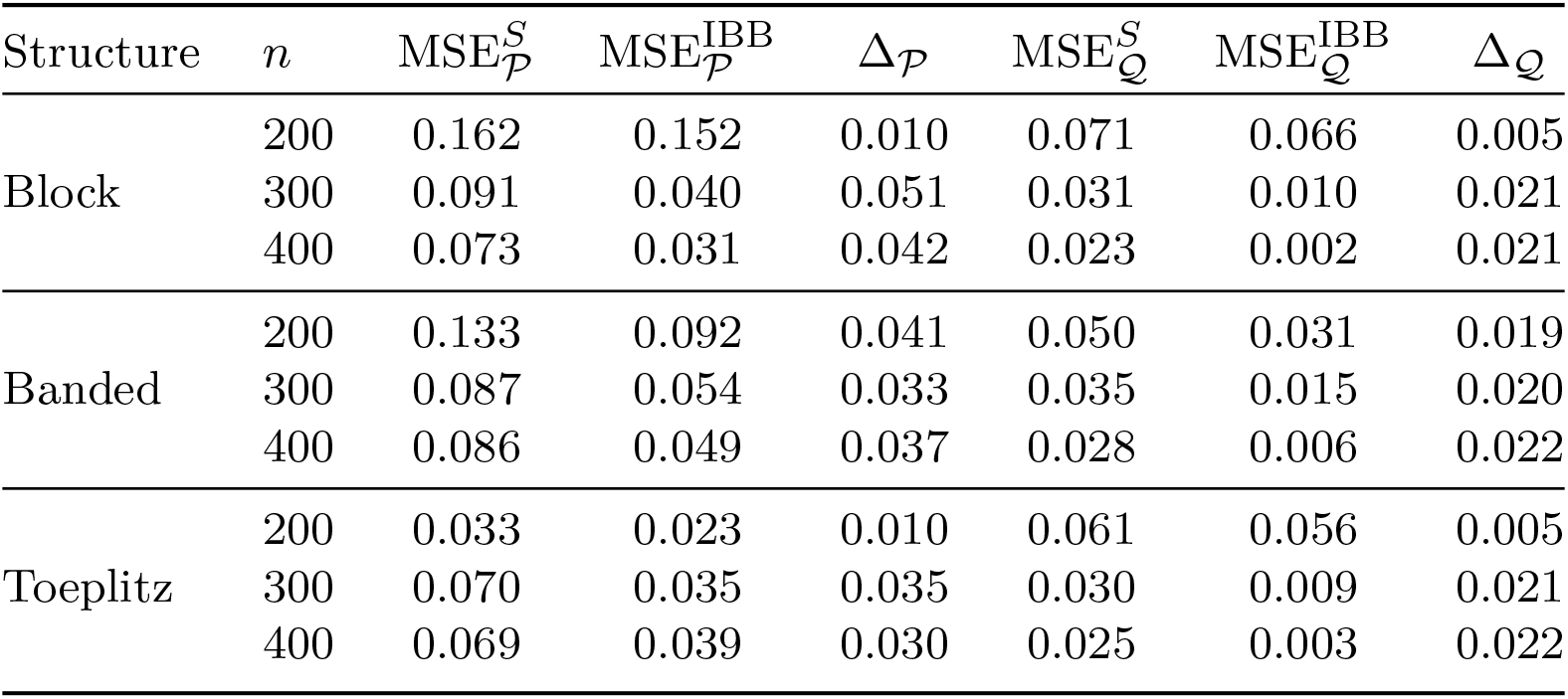
Element-wise mean squared errors on the correction-relevant subset *P* and its complement *Q* under the ultra-high-dimensional design *d* = 15 × *n* with *n* ∈ {200, 300, 400}.

**Table S6.**
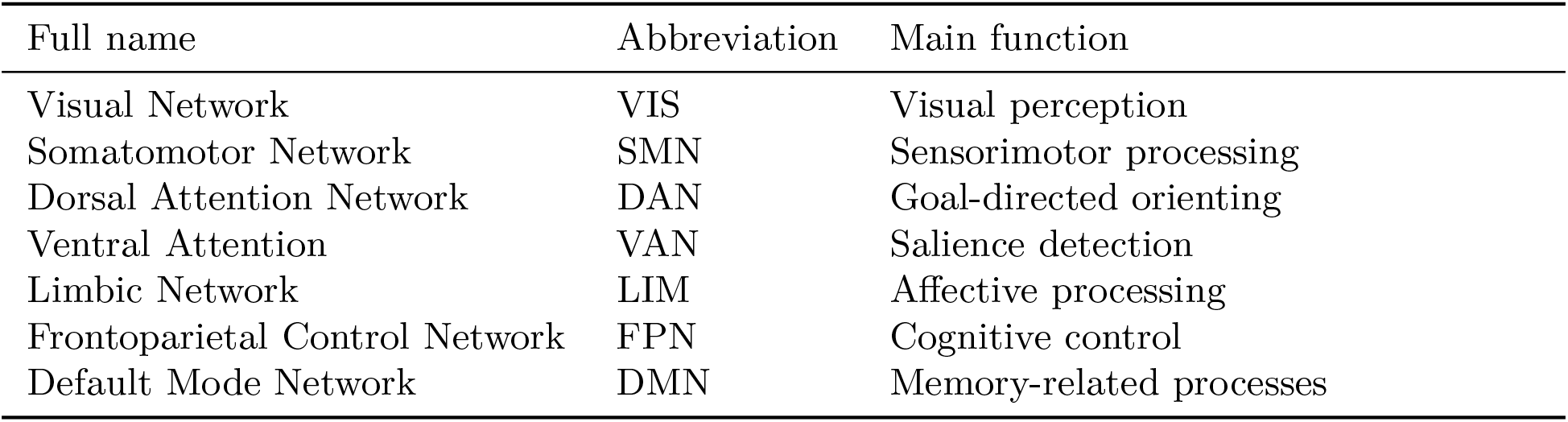
The seven canonical functional networks in the Yeo 7-network atlas.

**Table S7.**
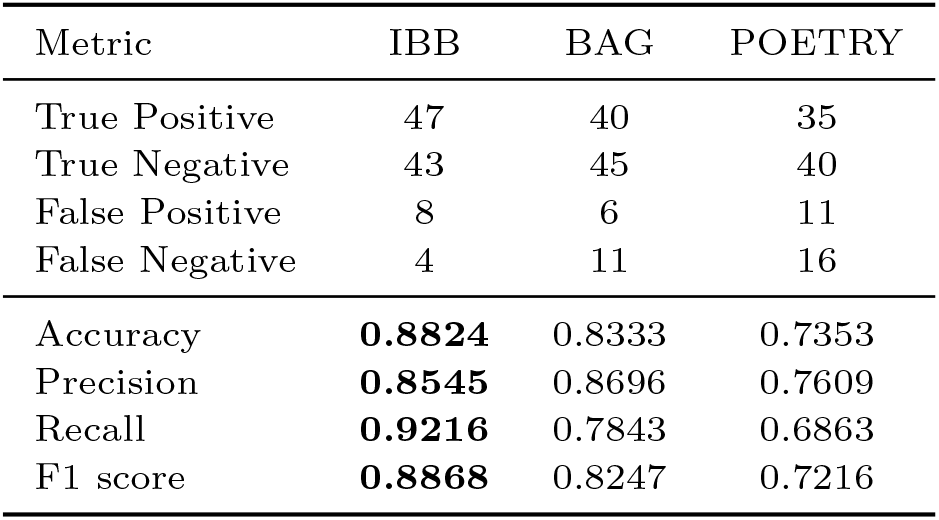
External validation performance on the NYU dataset.

### Supplementary Notes

#### Supplementary Note 1. Proofs of the theoretical results

In this section, we provide the proofs of the theoretical results stated in the main text. For clarity, each theorem, lemma, or remark is restated before its proof.

##### Theorem 1

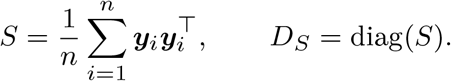

*Assume that the diagonal entries of S are strictly positive, so that* 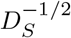 *is well defined. Let*

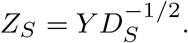

*For any* 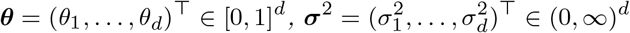, *and ε >* 0, *define*

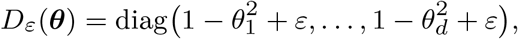

*and*

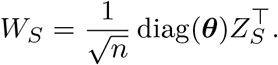

*Then the matrix*

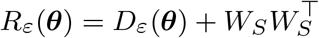

*is positive definite. Consequently, with*

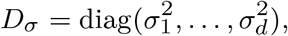

*the matrix*

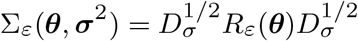

*is also positive definite*.

*Proof* For any nonzero vector ***x*** ∈ ℝ^*d*^,

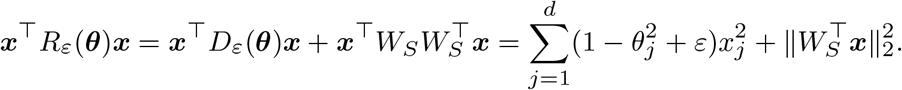

Since *θ*_*j*_ ∈ [0, 1] for every *j*, we have 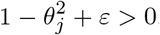. Hence the first term is strictly positive for all ***x*** ≠ **0**, while the second term is nonnegative. Therefore *R*_*ε*_(***θ***) is positive definite.

Moreover, since 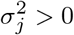 for every *j*, the matrix 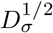 is nonsingular. Positive definiteness is preserved under congruence transformations by nonsingular matrices. Hence

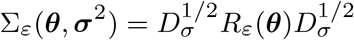

is positive definite.

##### Lemma 2

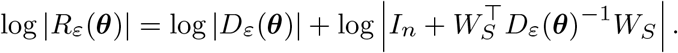

*Equivalently*,

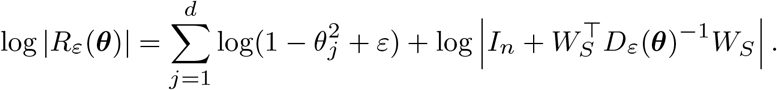

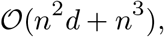

*which is dominated by O*(*n*^2^*d*) *whenever n*^3^ *is negligible relative to n*^2^*d*.
*Proof* Since

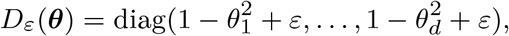

and, under the assumptions of Theorem 1, its diagonal entries are strictly positive, *D*_*ε*_(***θ***) is nonsingular. Applying the matrix determinant lemma to

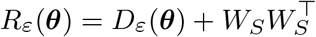

yields

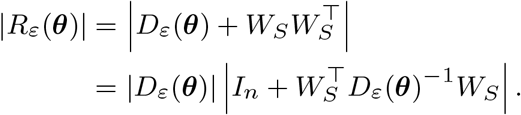

Taking logarithms gives

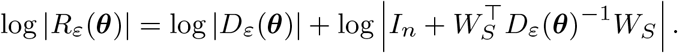

Because *D*_*ε*_(***θ***) is diagonal, we also have

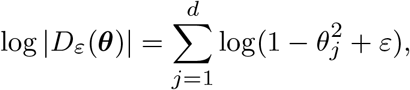

which proves the stated decomposition.

Next, we discuss the computational cost. A direct evaluation of

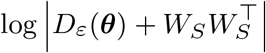

requires computing the log-determinant of a *d* × *d* matrix.

Using, for example, a Cholesky factorization, this step has the naive computational cost *O*(*d*^3^), in addition to the cost of forming the matrix itself. The latter consists of *<u>O</u>*(*d*) operations for computing the diagonal entries of *D*_*ε*_(***θ***) and *O* (*d^2^ n*) operations for forming 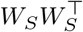. Thus, a direct evaluation incurs a total cost of order

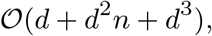

which, in high-dimensional settings where *n < d*, is typically dominated by *O*(*d*^3^).

By contrast, the decomposition above only requires the determinant of the *n* × *n* matrix

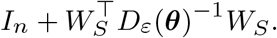

Forming

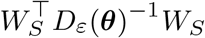

is equivalently to forming

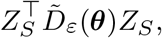

Where 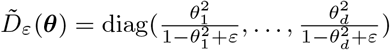. Since 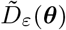 is diagonal, forming 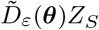 costs *O*(*dn*) operations only, thereby forming 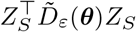 costs *O*(*n*^2^*d*) operations, subsequently computing the summation of *I*_*n*_ and the resulting *n* × *n* matrix, 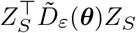, requires *O*(*n*) operations only as *I*_*n*_ is diagonal.

In addition, computing the log-determinant of the resulting *n*×*n* matrix, 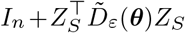, costs *O*(*n*^3^) operations via Cholesky factorization and the diagonal term

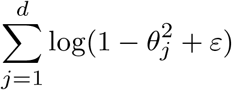

costs only *O*(*d*) operations.

Therefore, the total cost of the decomposed evaluation is

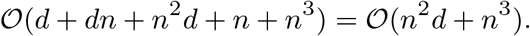

When *n* ≪ *d*, this is substantially smaller than the naive *O*(*d*^3^) cost. In particular, if *n*^3^ = *o*(*n*^2^*d*), then the leading order is *O*(*n*^2^*d*).

##### Lemma 3

(Trace decomposition) *Under the notation and assumptions of Theorem 1, the trace term satisfies*

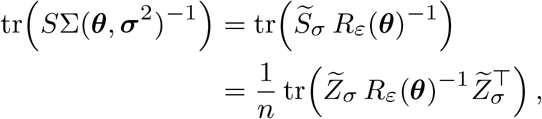

*Where* 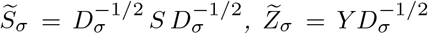. *Moreover, when n* ≪ *d, this decomposition reduces the computational cost of evaluating the trace term from the naive order O*(*d*^3^) *to*

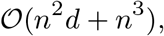

*which is dominated by O*(*n*^2^*d*) *whenever n*^3^ *is negligible relative to n*^2^*d*.

*Proof* According to the commutativity of trace operation with the decomposition of Σ(***θ, σ***^2^), we have

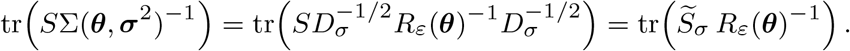

Where 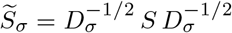.

Under the notation and assumptions of Theorem 1, the matrix 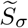 satisfies

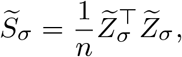

where 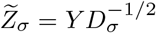. Thus we obtain the compressed form:

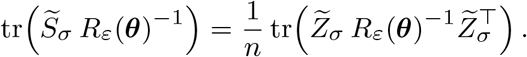

which proves the stated formula.

We next analyze the computational cost. Computing the inverse of Σ(***θ, σ***^2^)^−1^ has the naive computational cost *O*(*d*^3^), while forming a matrix as *S*Σ(***θ, σ***^2^)^−1^ costs *O*(*d*^3^). In addition, the trace computing is an *O*(*d*^2^) operation. Thus, a direct evaluation incurs a total cost of order

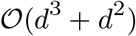

which is dominated by *O*(*d*^3^). By contrast, similar to Lemma 1, we apply the Woodbury identity to

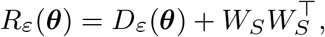

which yields an inverse expressed through the diagonal matrix *D*_*ε*_(***θ***)^−1^ and an *n* × *n* matrix

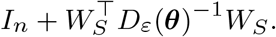

where *D*_*ε*_(***θ***)^−1^ costs *O*(*d*) up to forming diagonal entries, and forming 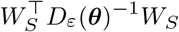 costs *O*(*n d*). In addition, computing the *n* × *n* inverse costs *O* (*n*) followed by the *O* (*n*) trace operation. Hence the total cost of the trace term is

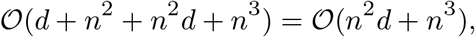

When *n* ≪ *d*, this is substantially smaller than the naive *O*(*d*^3^) cost. In particular, if *n*^3^ = *o*(*n*^2^*d*), then the leading order is *O*(*n*^2^*d*).

##### Theorem 4

(Conditioning bounds via Woodbury) *Under the notation and assumptions of Theorem 1, let 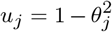 for j* = 1, …, *d. For an arbitrary real square matrix A, let λ*_min_(*A*) *and λ*_max_(*A*) *denote its smallest and largest eigenvalues, respectively*.

*Then*

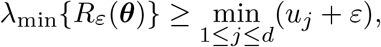

*and*

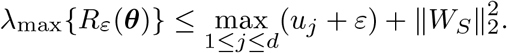

*Consequently, the condition number satisfies*

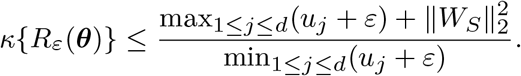

*Proof* We establish the lower and upper eigenvalue bounds separately.

We first prove the lower bound. For any ***ν*** ∈ ℝ^*d*^, we have

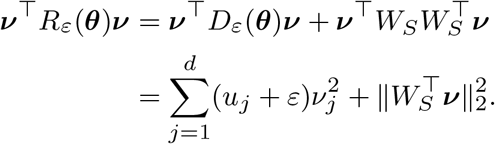

Since 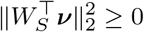, it follows that

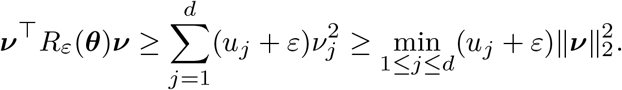

Moreover, *R*_*ε*_(***θ***) is real symmetric. Therefore, by the Rayleigh–Ritz characterization,

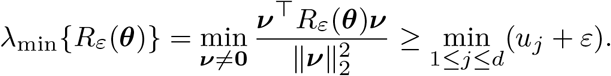

We next prove the upper bound. By Weyl’s inequality,

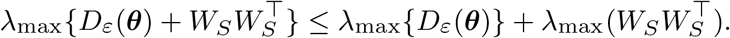

Since *D*_*ε*_(***θ***) is diagonal,

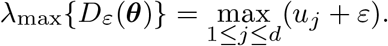

Furthermore, using the standard relationship between the spectral norm and the largest eigenvalue,

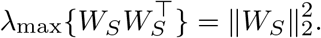

Consequently,

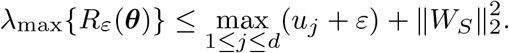

Combining the lower and upper bounds gives

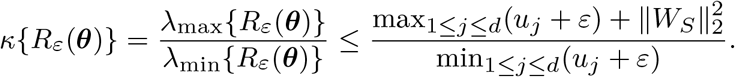

This completes the proof.

##### Theorem 5

(Posterior propriety) *Under the notation and assumptions of Theorem 1, suppose that the probability model is specified as in Eq. 1 (in Section Method). As described in Section Method, assume that the calibration parameters θ*_*j*_ *are determined by the global shrinkage parameter τ and the local shrinkage parameters λ*_*j*_, *j* = 1, …, *d, with the latter assigned proper half-Cauchy priors. Further assume that the variance components satisfy*

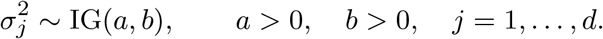

*Then, for any fixed ε >* 0, *the marginal normalizing constant*

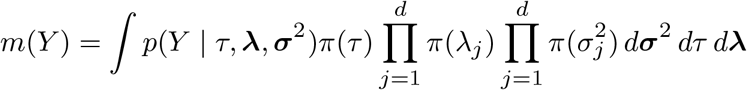

*is finite. Consequently, the posterior distribution is proper*.

*Proof* For any *ε >* 0, Theorem 4 implies that

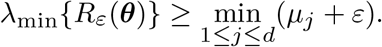

As stated in Theorem 4, we have *µ*_*j*_ = 1 − *θ*_*j*_ ≥ 0 for all *j* = 1, …, *d*. Therefore,

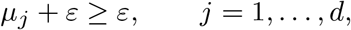

and hence

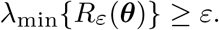

Consequently

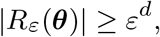

and therefore,

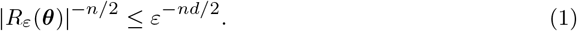

According to the probability model and the proposed parametrized structure for Σ(***θ***), the likelihood can be written as

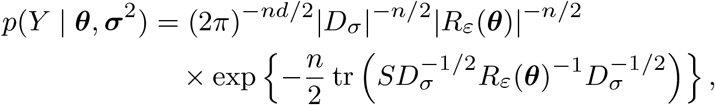

where

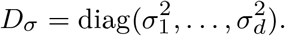

Next, we bound the exponential term. Define

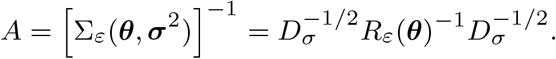

By Theorem 1, Σ_*ε*_(***θ, σ***^2^) is positive definite. Therefore, by standard properties of positive definite matrices, *A* is positive definite, and so is its principal square root *A*^1*/*2^.

Moreover, since *S* is positive semidefinite and positive semidefiniteness is preserved under congruence transformations by nonsingular matrices, it follows that

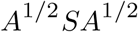

is positive semidefinite. Moreover, by the cyclic invariance of the trace,

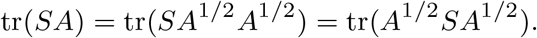

Since the trace of a positive semidefinite matrix is nonnegative, we obtain

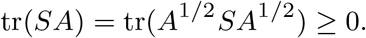

Therefore,

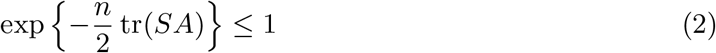

Combining the upper bound for the exponential term in Eq. (2) with the determinant bound in Eq. (1), we obtain

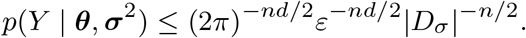

Let

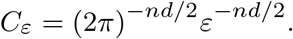

For any fixed *ε >* 0, we have *C*_*ε*_ *<* ∞.

Since

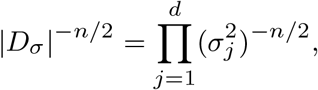

it follows that

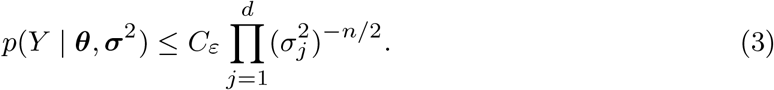

Substituting Eq. (3) into the marginal normalizing constant yields

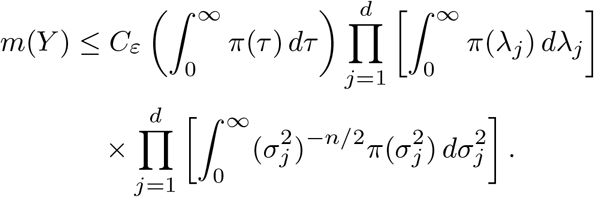

The half-Cauchy priors assigned to *τ* and *λ*_*j*_ are proper. Therefore,

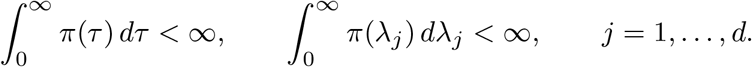

It remains to verify that each variance-related integral is finite. Since

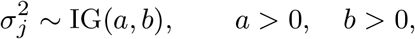

the density of 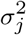 is

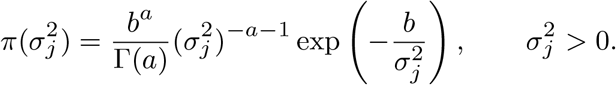

Hence,

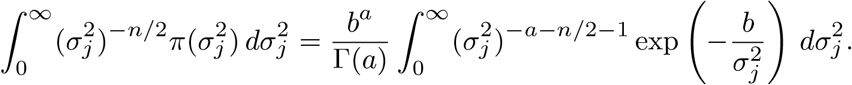

Using the change of variables

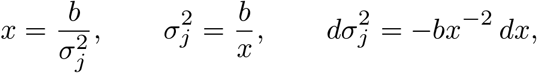

we obtain

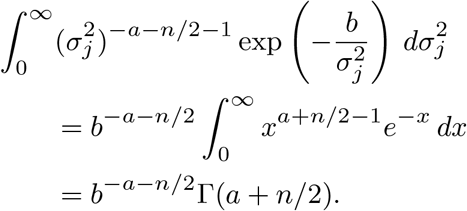

Consequently,

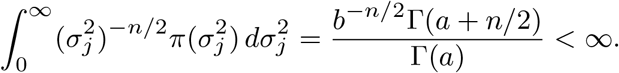

Since all factors in the upper bound for *m*(*Y*) are finite, we conclude that

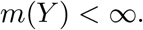

Therefore, the posterior distribution is proper.

##### Theorem 6

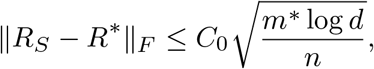

*where m*\* *denotes the number of nonzero off-diagonal entries in R*\*.

*Define the oracle approximation error*

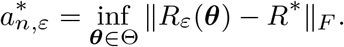

*Assume that* 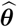 *satisfies the approximate projection condition*

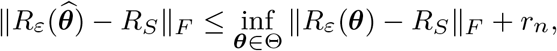

*where r*_*n*_ ≥ 0 *is the optimization or approximation tolerance. Then, on E*_*n*_,

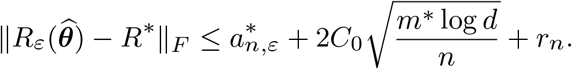

*In particular, if the support of R*\* *is induced by at most s active nodes, then*

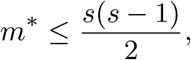

*and hence*

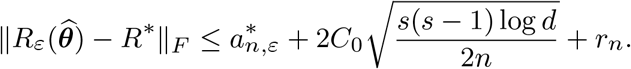

*Proof* We decompose the estimation error into an approximation component, a stochastic component, and an optimization tolerance component.

By the triangle inequality,

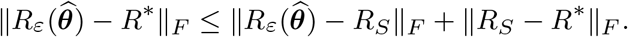

Using the approximate projection condition, we obtain

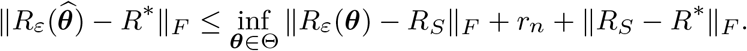

It remains to control the empirical projection term. For any ***θ*** ∈ Θ, another application of the triangle inequality gives

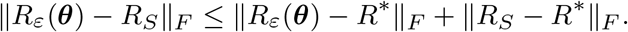

Taking the infimum over ***θ*** ∈ Θ on both sides yields

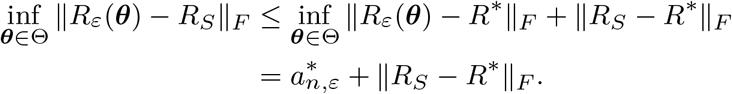

Combining the preceding inequalities gives

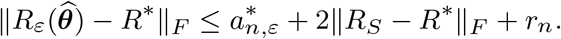

On the event *E*_*n*_, by assumption,

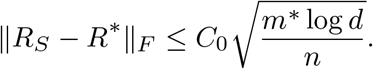

Therefore,

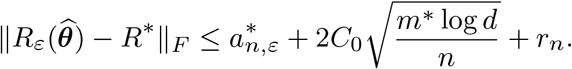

Finally, if the support of *R*\* is induced by at most *s* active nodes, then the maximum number of possible nonzero off-diagonal entries among these active nodes is

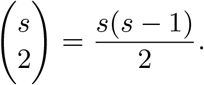

Thus,

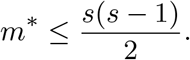

Substituting this bound into the preceding inequality gives

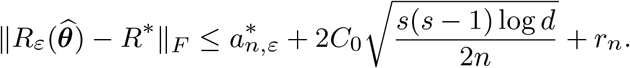

This completes the proof.

##### Theorem 7

*Under the notation and assumptions of Theorem 6, define*

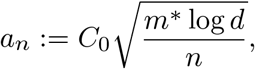

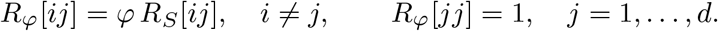

*Assume that the parameter φ >* 0 *satisfies*

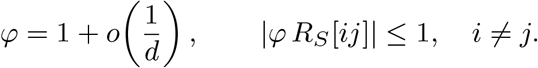

*Suppose that*

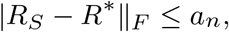

*and that the construction of R*_*φ*_ *yields the perturbation bound*

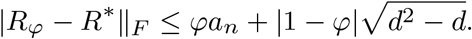

*Then, for all sufficiently large d, there exists a constant K >* 0 *such that*

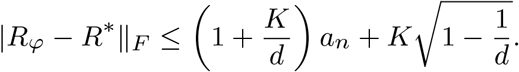

*More precisely, let*

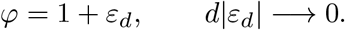

*Then*

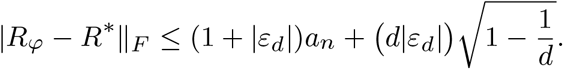

*Consequently*,

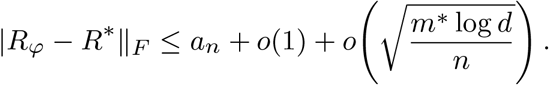

*In particular, if*

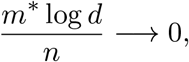

*then*

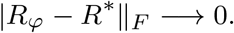

*Proof* Let

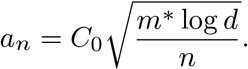

By the assumed perturbation bound,

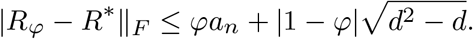

Substituting the definition of *a*_*n*_ gives

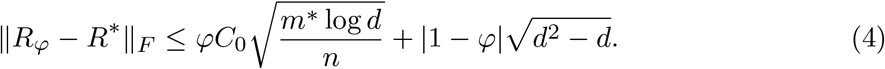

Since *φ* = 1 + *o*(1*/d*), there exists a sequence {*ε*_*d*_} such that

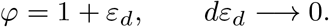

Thus |1 − *φ*| = |*ε*_*d*_|, and Eq. 4 becomes

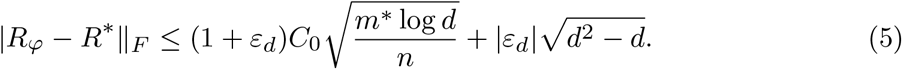

Noting that 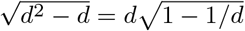, we have

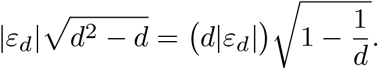

Using 1 + *ε*_*d*_ ≤ 1 + |*ε*_*d*_|, it follows from Eq. 5 that

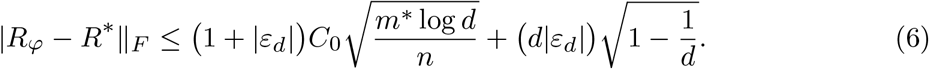

Because *d*|*ε*_*d*_| → 0, the sequence {*d*|*ε*_*d*_|} is bounded. Hence there exists a constant *K >* 0 such that, for all sufficiently large *d*,

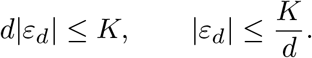

Substituting these into Eq. 6 yields

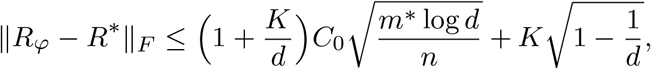

which establishes the finite-dimensional bound.

To derive the sharper asymptotic bound, expand Eq. 6 as

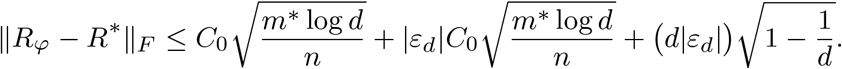

Since |*ε*_*d*_| = *o*(1*/d*) and *d*|*ε*_*d*_| = *o*(1), we obtain

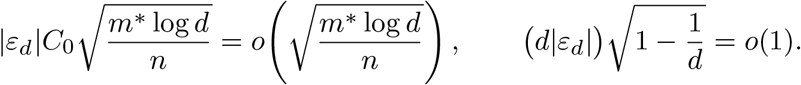

Therefore,

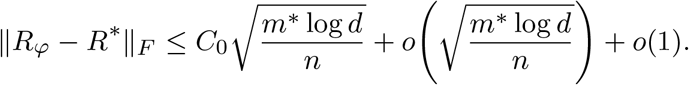

Finally, if (*m*\* log *d*)*/n* → 0, then 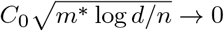, and the preceding bound gives

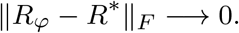

##### Remark 1

The Frobenius-norm error bound consists of two components:

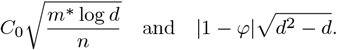

Since 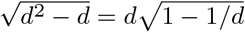, the perturbation term can be written as

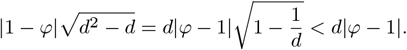

If this condition is not satisfied but the weaker condition 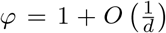 holds, then *d*|*φ* − 1| = *O*(1). In this case, the cumulative perturbation remains bounded.

#### Supplementary Note 2. The details of the covariance structures and the four MSE measures

This section covers the details of the covariance structures and the four MSE measures used in simulation studies.

##### Supplementary Note 2.1. Details of covariance structures

Here, we provide details of the three covariance structures—block, banded, and Toeplitz—used in the simulation studies, following [1–3].

1. **Block**. The indices 1, 2, …, *d* were partitioned into five ordered groups of equal size. Correlations were set to be moderate within each group and weak across groups, thereby mimicking clustered dependence patterns frequently encountered in brain connectivity studies:

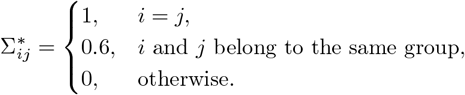
2. **Banded**. Two variables indexed by *i* and *j* were conditionally connected when |*i* − *j*| ≤ 3, yielding the covariance structure

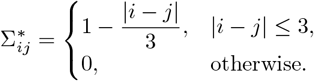
3. **Toeplitz**. Under the Toeplitz structure, covariance entries depended only on |*i* − *j* |, so that all elements along the same diagonal were equal:

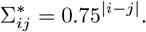

##### Supplementary Note 2.2. Four MSE measures

Four MSE measures were also used in the simulation studies to assess the effectiveness of the shrinkage terms and were calculated as belows, following [4]:

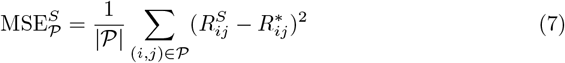

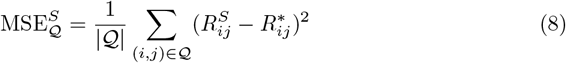

and

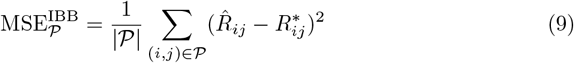

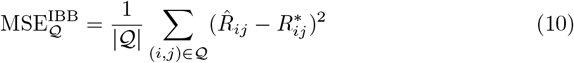

where 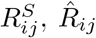 and 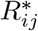 denote the entry of the sample correlation, posterior mean correlation and underlying true correlation matrix, respectively and

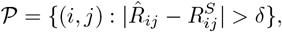

denotes the index set of entries whose absolute differences exceed a given threshold *δ*, and *Q* denotes the complement of *P*. We set *δ* = 0.05 to represent a practically meaningful level of correction [5].

#### Supplementary Note 3. Gaussian-likelihood-based classification

To assess downstream utility in the semi-synthetic study, we further evaluated a Gaussian-likelihood-based classification task. For each repetition, we performed 3-fold cross-validation on the semi-synthetic dataset. In each split, we used the training folds (with known group labels) to fit two Gaussian models, one for each group, and estimated the corresponding covariance matrices. Because our primary interest was covariance estimation, we set the group mean vectors 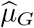 and 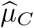 to the sample means of the training folds to simplify computation. We set the class prior to the empirical class proportion in the training fold, 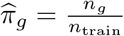. Similar to [6], for a validation observation **y**, we computed an unnormalized log-posterior score for each group:

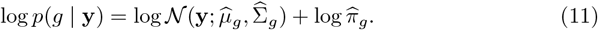

We then assigned **y** to the group with the larger score.

#### Supplementary Note 4. Validation using the New York University Child Study Center dataset

##### Supplementary Note 4.1. Dataset and preprocessing

We further evaluated the proposed method using the NYU ADHD-200 dataset. This validation analysis was designed to assess whether the group-level covariance structures learned from PKU dataset generalize to an independent ADHD cohort.

The original NYU sample exhibited substantial sex imbalance, with 51*/*48 male/female participants in the TDC group and 97*/*25 male/female participants in the ADHD group. Given this imbalance and prior evidence of sex-related differences in brain connectivity patterns, consistent with the motivation for the analysis of the PKU dataset, we restricted the validation analysis to male participants. To balance group sizes, male controls were randomly subsampled. The final analytical sample consisted of 51 TDCs and 51 ADHD cases, with each participant represented by features from 955 ROIs.

##### Supplementary Note 4.2. External validation based on covariance structures

To assess external validity, we used the PKU dataset as the training sample and the NYU dataset as an independent test sample. Specifically, we separately estimated the group-level covariance matrices for the ADHD and TDC groups from the PKU data, and then used these estimated covariance structures to classify participants in the NYU dataset following the same procedure as listed in Supplementary Note 3. The rationale of this analysis is that, if the covariance patterns learned from PKU remain effective for classifying subjects in NYU, then the group-level dependence structures underlying the two datasets are likely to be broadly consistent.

The classification results are summarized in Table S7. Among the three methods considered, the IBB achieved the best overall performance, with an accuracy of 0.8824, based on 47 true positives and 43 true negatives. The corresponding precision, recall, and F1 score were 0.8545, 0.9216, and 0.8868, respectively. Although the BAG attained a slightly higher precision (0.8696), the IBB outperformed the BAG and POETRY in terms of accuracy, recall, and F1 score, while the POETRY showed the weakest performance overall. Since all three methods were evaluated under the same PKU-training/NYU-testing framework, the superior classification performance of the IBB suggests that it provides a more accurate recovery of the case- and control-specific covariance structures. Taken together, these findings indicate that the covariance patterns estimated from the PKU data are reproducible in the NYU cohort and that the IBB captures these group-specific covariance differences more effectively than the BAG and POETRY.

##### Supplementary Note 4.3. Network-level differential analysis

We next examined whether the NYU dataset exhibited network-level group differences consistent with those observed in the main analysis. Each ROI was mapped to one of the canonical large-scale functional systems in the Yeo 7-network atlas [7–9]. Following the same block-level comparison procedure used for the PKU dataset, we summarized group differences between ADHD cases and TDCs at the level of network pairs.

Figure S1 presents the posterior mean correlation matrices for the TDC and ADHD groups, the relative difference heatmaps comparing the PKU and NYU mean correlation matrices for the two groups, and the block-wise permutation-based −log(*p*) and BH adjusted −log(*q*) heatmaps. For the relative difference panels, smaller values indicate greater cross-dataset agreement in network-level mean correlations. For the significance heatmaps, larger values indicate stronger evidence against the null hypothesis of no group difference. The conventional significance threshold of 0.05 corresponds to −log(0.05) = 2.995. After FDR correction, the network blocks involving DMN–VIS and SMN–LIM remained significant.

Overall, the NYU validation analysis yielded neurobiologically plausible group differences based on correlations of ROI-derived features. These findings are broadly consistent with the results obtained on the PKU dataset in the main application section, supporting the robustness and generalizability of the proposed covariance-based framework.

---

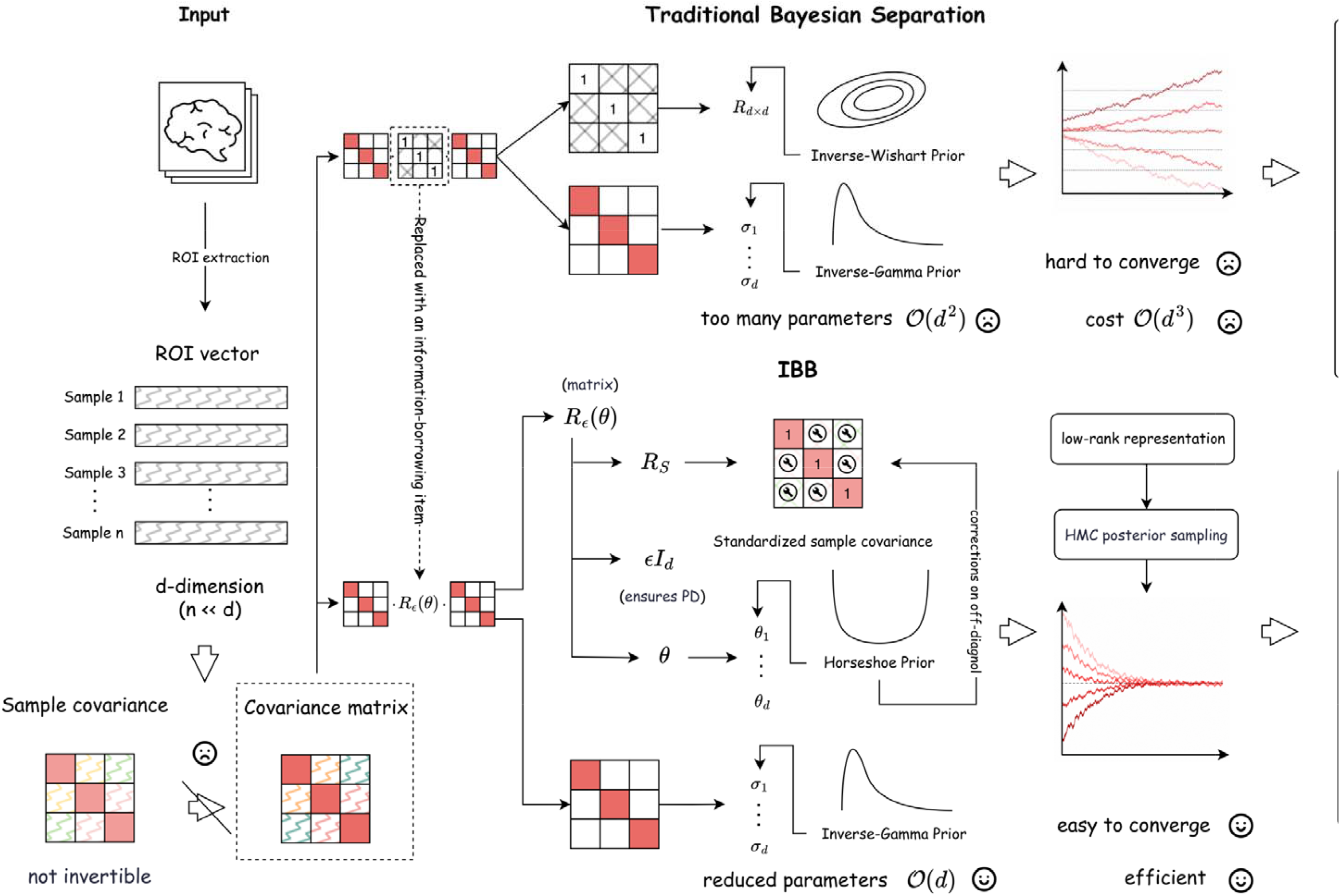

## References

[1] Hoff, G. A.-J., Van den Heuvel, M., Benders, M. J., Kersbergen, K. J. & De Vries, L. S. On development of functional brain connectivity in the young brain. Frontiers in human neuroscience 7, 650 (2013).

[2] Hagmann, P. et al. From diffusion mri to brain connectomics. (2005).

[3] Sporns, O., Tononi, G. & Köotter, R. The human connectome: a structural description of the human brain. PLoS computational biology 1, e42 (2005).

[4] Whitford, T. J., Kubicki, M. & Shenton, M. E. Diffusion tensor imaging, structural connectivity, and schizophrenia. Schizophrenia research and treatment 2011, 709523 (2011).

[5] Cabral, J. et al. Structural connectivity in schizophrenia and its impact on the dynamics of spontaneous functional networks. Chaos: An Interdisciplinary Journal of Nonlinear Science 23 (2013).

[6] Wheeler, A. L. & Voineskos, A. N. A review of structural neuroimaging in schizophrenia: from connectivity to connectomics. Frontiers in human neuroscience 8, 653 (2014).

[7] He, B. & Liu, Z. Multimodal functional neuroimaging: integrating functional mri and eeg/meg. IEEE reviews in biomedical engineering 1, 23–40 (2008).

[8] Babiloni, C., Pizzella, V., Del Gratta, C., Ferretti, A. & Romani, G. L. Fundamentals of electroencefalography, magnetoencefalography, and functional magnetic resonance imaging. International review of neurobiology 86, 67–80 (2009).

[9] Baillet, S. Magnetoencephalography for brain electrophysiology and imaging. Nature neuroscience 20, 327–339 (2017).

[10] Zalesky, A. et al. Whole-brain anatomical networks: does the choice of nodes matter? Neuroimage 50, 970–983 (2010).

[11] Kaiser, M. A tutorial in connectome analysis: topological and spatial features of brain networks. Neuroimage 57, 892–907 (2011).

[12] Stanley, M. L. et al. Defining nodes in complex brain networks. Frontiers in computational neuroscience 7, 169 (2013).

[13] Huang, S. et al. Learning brain connectivity of alzheimer’s disease by sparse inverse covariance estimation. NeuroImage 50, 935–949 (2010).

[14] Zhou, L., Wang, L. & Ogunbona, P. Discriminative sparse inverse covariance matrix: application in brain functional network classification. Proceedings of the IEEE Conference on Computer Vision and Pattern Recognition 39, 3097–3104 (2014).

[15] Dai, M., Zhang, Z. & Srivastava, A. Analyzing dynamical brain functional connectivity as trajectories on space of covariance matrices. IEEE transactions on medical imaging 39, 611–620 (2019).

[16] Bullmore, E. & Sporns, O. Complex brain networks: graph theoretical analysis of structural and functional systems. Nature reviews neuroscience 10, 186–198 (2009).

[17] Friston, K. J. Functional and effective connectivity: a review. Brain connectivity 1, 13–36 (2011).

[18] Lam, C. High-dimensional covariance matrix estimation. Wiley Interdisciplinary reviews: computational statistics 12, e1485 (2020).

[19] Fan, J., Liao, Y. & Mincheva, M. Large covariance estimation by thresholding principal orthogonal complements. Journal of the Royal Statistical Society Series B: Statistical Methodology 75, 603–680 (2013).

[20] Zou, S. & Zhao, Z. Large covariance matrix estimation based on factor models via nonconvex optimization 1, 9656–9660 (2024).

[21] Barnard, J., McCulloch, R. & Meng, X.-L. Modeling covariance matrices in terms of standard deviations and correlations, with application to shrinkage. Statistica Sinica 1281–1311 (2000).

[22] Lan, S. et al. Flexible bayesian dynamic modeling of correlation and covariance matrices. Bayesian analysis 15, 1199 (2019).

[23] Wang, C., Du, J. & Fan, X. High-dimensional correlation matrix estimation for general continuous data with bagging technique. Machine Learning 111, 2905–2927 (2022).

[24] Tipping, M. E. & Bishop, C. M. Probabilistic principal component analysis. Journal of the Royal Statistical Society Series B: Statistical Methodology 61, 611–622 (1999).

[25] Bellec, P. et al. The neuro bureau adhd-200 preprocessed repository. Neuroimage 144, 275–286 (2017).

[26] Thomas Yeo, B. et al. The organization of the human cerebral cortex estimated by intrinsic functional connectivity. Journal of neurophysiology 106, 1125–1165 (2011).

[27] Kay, B. P. et al. Stimulant medications affect arousal and reward, not attention networks. Cell 188, 7529–7546 (2025).

[28] Yeo, B. T. et al. The organization of the human cerebral cortex estimated by intrinsic functional connectivity. Journal of neurophysiology (2011).

[29] Benjamini, Y. & Hochberg, Y. Controlling the false discovery rate: a practical and powerful approach to multiple testing. Journal of the Royal statistical society: series B (Methodological) 57, 289–300 (1995).

[30] Smallwood, J. et al. The default modes of reading: modulation of posterior cingulate and medial prefrontal cortex connectivity associated with comprehension and task focus while reading. Frontiers in human neuroscience 7, 734 (2013).

[31] Zhang, M. et al. Perceptual coupling and decoupling of the default mode network during mind-wandering and reading. elife 11, e74011 (2022).

[32] Xu, S. et al. Functional connectivity profiles of the default mode and visual networks reflect temporal accumulative effects of sustained naturalistic emotional experience. NeuroImage 269, 119941 (2023).

[33] Fernandino, L. & Binder, J. R. How does the “default mode” network contribute to semantic cognition? Brain and language 252, 105405 (2024).

[34] Rolls, E. T. The cingulate cortex and limbic systems for emotion, action, and memory. Brain structure and function 224, 3001–3018 (2019).

[35] Braine, A. & Georges, F. Emotion in action: When emotions meet motor circuits. Neuroscience & Biobehavioral Reviews 155, 105475 (2023).

[36] Fan, J. & Lv, J. Sure independence screening for ultrahigh dimensional feature space. Journal of the Royal Statistical Society Series B: Statistical Methodology 70, 849–911 (2008).

[37] Chen, T., Fox, E. & Guestrin, C. Stochastic gradient hamiltonian monte carlo 1683–1691 (2014).

[38] Neal, R. M. et al. Mcmc using hamiltonian dynamics. Handbook of markov chain monte carlo 2, 2 (2011).

[39] Betancourt, M. A conceptual introduction to hamiltonian monte carlo. arXiv preprint arXiv:1701.02434 (2017).

[40] Harville, D. A. Matrix algebra from a statistician’s perspective (1998).

[41] Rothman, A. J., Levina, E. & Zhu, J. Generalized thresholding of large covariance matrices. Journal of the American Statistical Association 104, 177–186 (2009).

[42] Xue, L., Ma, S. & Zou, H. Positive-definite l1-penalized estimation of large covariance matrices. Journal of the American Statistical Association 107, 1480–1491 (2012).

[43] Cui, Y., Leng, C. & Sun, D. Sparse estimation of high-dimensional correlation matrices. Computational Statistics & Data Analysis 93, 390–403 (2016).

[44] Jacobson, L. A. et al. Sex-based dissociation of white matter microstructure in children with attention-deficit/hyperactivity disorder. Journal of the American Academy of Child & Adolescent Psychiatry 54, 938–946 (2015).

[45] Dupont, G., van Rooij, D., Buitelaar, J. K., Reif, A. & Grimm, O. Sex-related differences in adult attention-deficit hyperactivity disorder patients–an analysis of external globus pallidus functional connectivity in resting-state functional mri. Frontiers in psychiatry 13, 962911 (2022).

[46] Carucci, S. et al. Clinical characteristics, neuroimaging findings, and neuropsychological functioning in attention-deficit hyperactivity disorder: Sex differences. Journal of Neuroscience Research 101, 704–717 (2023).

[47] Winkler, A. M., Ridgway, G. R., Webster, M. A., Smith, S. M. & Nichols, T. E. Permutation inference for the general linear model. Neuroimage 92, 381–397 (2014).

[48] Phipson, B. & Smyth, G. K. Permutation p-values should never be zero: calculating exact p-values when permutations are randomly drawn. arXiv preprint arXiv:1603.05766 (2016).

[49] North, B. V., Curtis, D. & Sham, P. C. A note on the calculation of empirical p values from monte carlo procedures. The American Journal of Human Genetics 71, 439–441 (2002).

## References

[1] Rothman, A. J., Levina, E. & Zhu, J. Generalized thresholding of large covariance matrices. Journal of the American Statistical Association 104, 177–186 (2009).

[2] Xue, L., Ma, S. & Zou, H. Positive-definite l1-penalized estimation of large covariance matrices. Journal of the American Statistical Association 107, 1480–1491 (2012).

[3] Cui, Y., Leng, C. & Sun, D. Sparse estimation of high-dimensional correlation matrices. Computational Statistics & Data Analysis 93, 390–403 (2016).

[4] Ledoit, O. & Wolf, M. Improved estimation of the covariance matrix of stock returns with an application to portfolio selection. Journal of empirical finance 10, 603–621 (2003).

[5] Amrhein, V., Greenland, S. & McShane, B. Scientists rise up against statistical significance. Nature 567, 305–307 (2019).

[6] Lee, K., Jo, S., Lee, K. & Lee, J. Scalable and optimal bayesian inference for sparse covariance matrices via screened beta-mixture prior. Bayesian Analysis 1, 1–28 (2024).

[7] Thomas Yeo, B. et al. The organization of the human cerebral cortex estimated by intrinsic functional connectivity. Journal of neurophysiology 106, 1125–1165 (2011).

[8] Kay, B. P. et al. Stimulant medications affect arousal and reward, not attention networks. Cell 188, 7529–7546 (2025).

[9] Yeo, B. T. et al. The organization of the human cerebral cortex estimated by intrinsic functional connectivity. Journal of neurophysiology (2011).

