## Supplementary for "An information-borrowing Bayesian method for inferring high-dimensional population-level brain connectivity from small samples"

### Supplementary Information

### Supplementary Figures

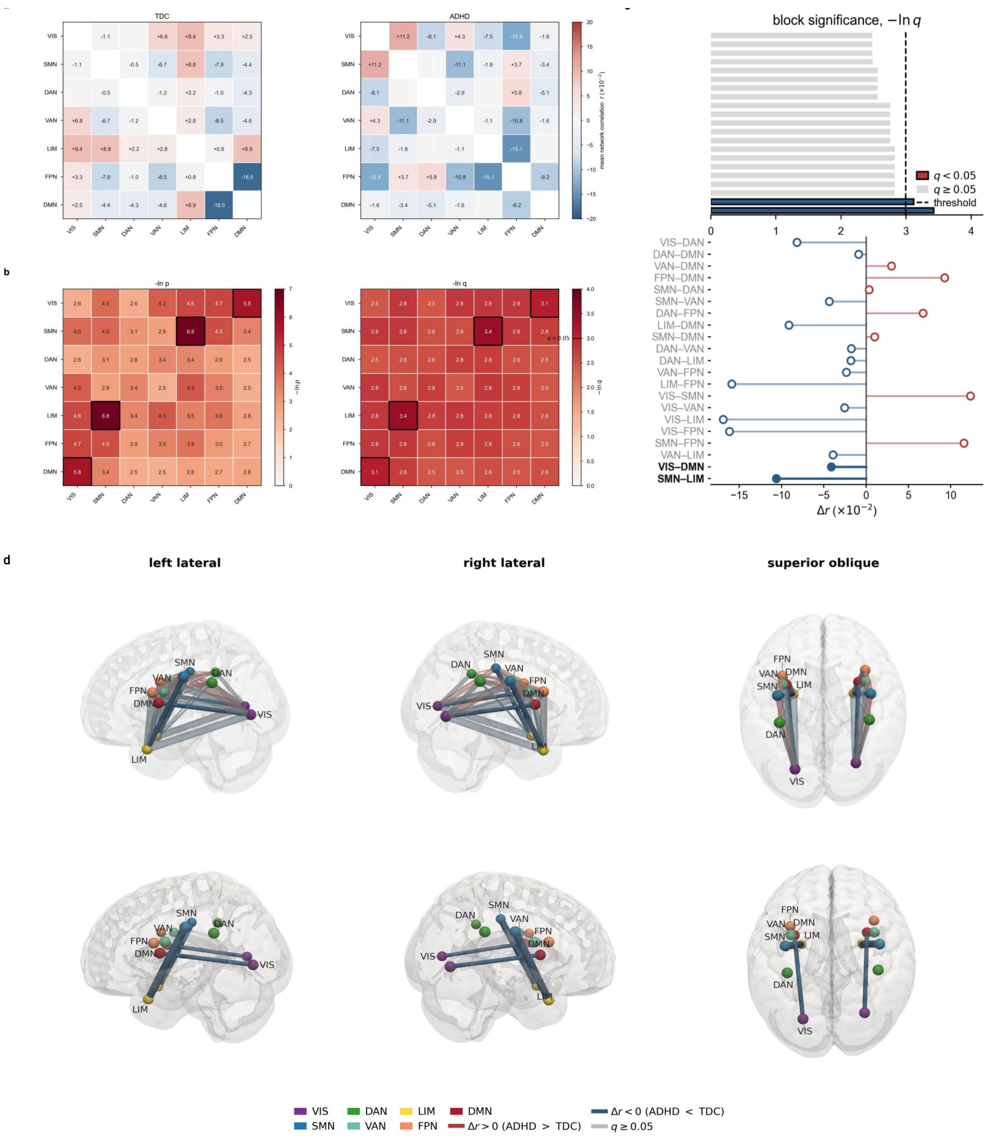

Fig. S1. Network-level group comparison in the NYU ADHD200 dataset. **a**, Posterior mean correlation matrix for the TDC group (left) and ADHD group (right). **b**, Heatmap of block-wise permutation test significance, shown as $-\log(p)$ (left) and Heatmap of BH-FDR adjusted significance, shown as $-\log(q)$ (right). Network blocks correspond to pairs of Yeo 7 functional systems, including VIS, SMN, DAN, VAN, LIM, FPN, and DMN. In this panel, larger values indicate stronger evidence of group differences between the TDC and ADHD group. **c**, Block-level effect size. The upper bars give the BH-FDR adjusted significance of each network pair, $-\log(q)$; the lower lollipops give the signed difference in posterior mean correlation, $\Delta r = r_{\text{ADHD}} - r_{\text{TDC}}$. Color encodes the sign of $\Delta r$ (red positive, blue negative). The two FDR-significant blocks are drawn in bold with their pair names emphasised. **d**, Spatial rendering of ADHD-minus-TDC differences in network-level posterior mean correlations. The cortical surface was derived from the MNI152NLin2009cAsym T1-weighted template after Gaussian smoothing ($\sigma = 1\,\text{mm}$), using an isosurface approximating the grey matter–cerebrospinal fluid boundary, and shaded according to mean curvature. Nodes represent the centroids of the seven Yeo networks within each hemisphere (14 nodes in total). Tube color encodes the signed group difference, $\Delta r$, and tube radius is proportional to $|\Delta r|$. Each displayed network-pair effect is rendered once in each hemisphere for visualization only; the two renderings represent the same block-level comparison, not separate hemisphere-specific estimates or tests. The second row displays only network pairs with Benjamini–Hochberg-adjusted $q < 0.05$.

### Supplementary Tables

Table S1. Mean estimation errors with standard deviations (in parentheses) over 100 replications when $n = 100$ and $d = 200$. Errors are reported in Frobenius norm and spectral norm.

| Structure | Measure | SCM | BAG | IBB | POETRY |
|---|---|---|---|---|---|
| | $\|\cdot\|_F$ | 20.604 | 20.565 | **19.271** | 19.448 |
| | | (1.585) | (1.544) | (1.324) | (1.535) |
| Block | $\|\cdot\|_2$ | 11.462 | 11.418 | **10.380** | 11.294 |
| | | (1.808) | (1.761) | (1.416) | (1.706) |
| | PD | 0/100 | **100/100** | **100/100** | **100/100** |
| | $\|\cdot\|_F$ | 20.372 | 20.337 | **18.672** | 20.235 |
| | | (0.757) | (0.758) | (0.545) | (0.738) |
| Banded | $\|\cdot\|_2$ | 8.465 | 8.431 | **6.914** | 8.336 |
| | | (0.807) | (0.804) | (0.620) | (0.798) |
| | PD | 0/100 | **100/100** | **100/100** | **100/100** |
| | $\|\cdot\|_F$ | 20.263 | 20.222 | **15.355** | 20.193 |
| | | (0.504) | (0.504) | (0.232) | (0.494) |
| Toeplitz | $\|\cdot\|_2$ | 7.050 | 7.035 | **4.465** | 6.974 |
| | | (0.531) | (0.529) | (0.195) | (0.525) |
| | PD | 0/100 | **100/100** | **100/100** | **100/100** |

Table S2. Mean estimation errors with standard deviations (in parentheses) over 100 replications when $n = 100$ and $d = 500$. Errors are reported in Frobenius norm and spectral norm.

| Structure | Measure | SCM | BAG | IBB | POETRY |
|---|---|---|---|---|---|
| Block | $\|\cdot\|_F$ | 50.931 | 50.806 | **49.478** | 50.052 |
| | | (4.042) | (4.031) | (3.542) | (3.934) |
| | $\|\cdot\|_2$ | 27.588 | 27.476 | **26.396** | 27.190 |
| | | (5.115) | (5.052) | (4.386) | (4.933) |
| | PD | 0/100 | **100/100** | **100/100** | **100/100** |
| Banded | $\|\cdot\|_F$ | 50.747 | 50.555 | **49.582** | 50.813 |
| | | (1.167) | (1.159) | (1.101) | (1.149) |
| | $\|\cdot\|_2$ | 16.066 | 15.995 | **15.514** | 15.904 |
| | | (1.162) | (1.158) | (1.129) | (1.150) |
| | PD | 0/100 | **100/100** | **100/100** | **100/100** |
| Toeplitz | $\|\cdot\|_F$ | 50.529 | 50.333 | **41.065** | 50.104 |
| | | (0.719) | (0.715) | (0.592) | (0.718) |
| | $\|\cdot\|_2$ | 13.483 | 13.422 | **10.077** | 13.448 |
| | | (0.791) | (0.781) | (0.633) | (0.781) |
| | PD | 0/100 | **100/100** | **100/100** | **100/100** |

Table S3. Mean estimation errors with standard deviations (in parentheses) over 100 replications when $n = 100$ and $d = 1 \times 10^3$. Errors are reported in Frobenius norm and spectral norm.

| Structure | Measure | SCM | BAG | IBB | POETRY |
|---|---|---|---|---|---|
| Block | $\|\cdot\|_F$ | 103.767 | 103.498 | **98.405** | 101.252 |
| | | (8.003) | (7.899) | (8.246) | (7.937) |
| | $\|\cdot\|_2$ | 57.416 | 56.286 | **55.437** | 56.101 |
| | | (8.699) | (8.532) | (7.148) | (8.302) |
| | PD | 0/100 | **100/100** | **100/100** | **100/100** |
| Banded | $\|\cdot\|_F$ | 100.688 | 100.270 | **90.583** | 100.188 |
| | | (1.204) | (1.208) | (0.993) | (1.208) |
| | $\|\cdot\|_2$ | 25.744 | 25.675 | **22.379** | 25.651 |
| | | (1.100) | (1.119) | (0.971) | (1.092) |
| | PD | 0/100 | **100/100** | **100/100** | **100/100** |
| Toeplitz | $\|\cdot\|_F$ | 100.582 | 100.156 | **89.669** | 91.428 |
| | | (0.843) | (0.845) | (0.713) | (0.820) |
| | $\|\cdot\|_2$ | 21.991 | 21.929 | **19.179** | 21.983 |
| | | (0.720) | (0.728) | (0.628) | (0.712) |
| | PD | 0/100 | **100/100** | **100/100** | **100/100** |

Table S4. Element-wise mean squared errors on the correction-relevant subset $\mathcal{P}$ and its complement $\mathcal{Q}$ under the moderate-dimensional design with fixed sample size $n = 100$ and dimensions $d \in \{200, 500, 1 \times 10^3\}$.

| Structure | $d$ | $\text{MSE}_{\mathcal{P}}^{S}$ | $\text{MSE}_{\mathcal{P}}^{\text{IBB}}$ | $\Delta_{\mathcal{P}}$ | $\text{MSE}_{\mathcal{Q}}^{S}$ | $\text{MSE}_{\mathcal{Q}}^{\text{IBB}}$ | $\Delta_{\mathcal{Q}}$ |
|---|---|---|---|---|---|---|---|
| | 200 | 0.045 | 0.040 | 0.005 | 0.020 | 0.017 | 0.003 |
| Block | 500 | 0.081 | 0.070 | 0.011 | 0.035 | 0.028 | 0.007 |
| | 1000 | 0.113 | 0.096 | 0.017 | 0.050 | 0.040 | 0.010 |
| | 200 | 0.052 | 0.046 | 0.006 | 0.025 | 0.021 | 0.004 |
| Banded | 500 | 0.086 | 0.073 | 0.013 | 0.038 | 0.030 | 0.008 |
| | 1000 | 0.121 | 0.101 | 0.020 | 0.055 | 0.044 | 0.011 |
| | 200 | 0.030 | 0.026 | 0.004 | 0.040 | 0.035 | 0.005 |
| Toeplitz | 500 | 0.068 | 0.060 | 0.008 | 0.058 | 0.050 | 0.008 |
| | 1000 | 0.095 | 0.082 | 0.013 | 0.072 | 0.060 | 0.012 |

Table S5. Element-wise mean squared errors on the correction-relevant subset $\mathcal{P}$ and its complement $\mathcal{Q}$ under the ultra-high-dimensional design $d = 15 \times n$ with $n \in \{200, 300, 400\}$.

| Structure | $n$ | $\text{MSE}_{\mathcal{P}}^{S}$ | $\text{MSE}_{\mathcal{P}}^{\text{IBB}}$ | $\Delta_{\mathcal{P}}$ | $\text{MSE}_{\mathcal{Q}}^{S}$ | $\text{MSE}_{\mathcal{Q}}^{\text{IBB}}$ | $\Delta_{\mathcal{Q}}$ |
|---|---|---|---|---|---|---|---|
| | 200 | 0.162 | 0.152 | 0.010 | 0.071 | 0.066 | 0.005 |
| Block | 300 | 0.091 | 0.040 | 0.051 | 0.031 | 0.010 | 0.021 |
| | 400 | 0.073 | 0.031 | 0.042 | 0.023 | 0.002 | 0.021 |
| | 200 | 0.133 | 0.092 | 0.041 | 0.050 | 0.031 | 0.019 |
| Banded | 300 | 0.087 | 0.054 | 0.033 | 0.035 | 0.015 | 0.020 |
| | 400 | 0.086 | 0.049 | 0.037 | 0.028 | 0.006 | 0.022 |
| | 200 | 0.033 | 0.023 | 0.010 | 0.061 | 0.056 | 0.005 |
| Toeplitz | 300 | 0.070 | 0.035 | 0.035 | 0.030 | 0.009 | 0.021 |
| | 400 | 0.069 | 0.039 | 0.030 | 0.025 | 0.003 | 0.022 |

Table S6. The seven canonical functional networks in the Yeo 7-network atlas.

| Full name | Abbreviation | Main function |
|---|---|---|
| Visual Network | VIS | Visual perception |
| Somatomotor Network | SMN | Sensorimotor processing |
| Dorsal Attention Network | DAN | Goal-directed orienting |
| Ventral Attention | VAN | Salience detection |
| Limbic Network | LIM | Affective processing |
| Frontoparietal Control Network | FPN | Cognitive control |
| Default Mode Network | DMN | Memory-related processes |

Table S7. External validation performance on the NYU dataset.

| Metric | IBB | BAG | POETRY |
|---|---|---|---|
| True Positive | 47 | 40 | 35 |
| True Negative | 43 | 45 | 40 |
| False Positive | 8 | 6 | 11 |
| False Negative | 4 | 11 | 16 |
| Accuracy | **0.8824** | 0.8333 | 0.7353 |
| Precision | **0.8545** | 0.8696 | 0.7609 |
| Recall | **0.9216** | 0.7843 | 0.6863 |
| F1 score | **0.8868** | 0.8247 | 0.7216 |

### Supplementary Notes

#### Supplementary Note 1. Proofs of the theoretical results

In this section, we provide the proofs of the theoretical results stated in the main text. For clarity, each theorem, lemma, or remark is restated before its proof.

**Theorem 1** *Let $Y = (y_{ij}) \in \mathbb{R}^{n\times d}$, with rows $\boldsymbol{y}_i = (y_{i1}, \dots, y_{id})^\top \in \mathbb{R}^d$, $i = 1, \dots, n$, and define*

$$S = \frac{1}{n}\sum_{i=1}^{n} \boldsymbol{y}_i \boldsymbol{y}_i^\top, \qquad D_S = \mathrm{diag}(S).$$

*Assume that the diagonal entries of $S$ are strictly positive, so that $D_S^{-1/2}$ is well defined. Let*

$$Z_S = Y D_S^{-1/2}.$$

*For any $\boldsymbol{\theta} = (\theta_1, \dots, \theta_d)^\top \in [0,1]^d$, $\boldsymbol{\sigma}^2 = (\sigma_1^2, \dots, \sigma_d^2)^\top \in (0, \infty)^d$, and $\varepsilon > 0$, define*

$$D_\varepsilon(\boldsymbol{\theta}) = \mathrm{diag}\big(1 - \theta_1^2 + \varepsilon, \dots, 1 - \theta_d^2 + \varepsilon\big),$$

*and*

$$W_S = \frac{1}{\sqrt{n}}\,\mathrm{diag}(\boldsymbol{\theta}) Z_S^\top.$$

*Then the matrix*

$$R_\varepsilon(\boldsymbol{\theta}) = D_\varepsilon(\boldsymbol{\theta}) + W_S W_S^\top$$

*is positive definite. Consequently, with*

$$D_\sigma = \mathrm{diag}(\sigma_1^2, \dots, \sigma_d^2),$$

*the matrix*

$$\Sigma_\varepsilon(\boldsymbol{\theta}, \boldsymbol{\sigma}^2) = D_\sigma^{1/2} R_\varepsilon(\boldsymbol{\theta}) D_\sigma^{1/2}$$

*is also positive definite.*

*Proof* For any nonzero vector $\boldsymbol{x} \in \mathbb{R}^d$,

$$\boldsymbol{x}^\top R_\varepsilon(\boldsymbol{\theta})\boldsymbol{x} = \boldsymbol{x}^\top D_\varepsilon(\boldsymbol{\theta})\boldsymbol{x} + \boldsymbol{x}^\top W_S W_S^\top \boldsymbol{x} = \sum_{j=1}^{d}(1-\theta_j^2+\varepsilon)x_j^2 + \|W_S^\top \boldsymbol{x}\|_2^2.$$

Since $\theta_j \in [0,1]$ for every $j$, we have $1-\theta_j^2+\varepsilon > 0$. Hence the first term is strictly positive for all $\boldsymbol{x} \neq \mathbf{0}$, while the second term is nonnegative. Therefore $R_\varepsilon(\boldsymbol{\theta})$ is positive definite.

Moreover, since $\sigma_j^2 > 0$ for every $j$, the matrix $D_\sigma^{1/2}$ is nonsingular. Positive definiteness is preserved under congruence transformations by nonsingular matrices. Hence

$$\Sigma_\varepsilon(\boldsymbol{\theta}, \boldsymbol{\sigma}^2) = D_\sigma^{1/2} R_\varepsilon(\boldsymbol{\theta}) D_\sigma^{1/2}$$

is positive definite. □

**Lemma 2** (Log-determinant decomposition) *Under the notation and assumptions of Theorem 1, the matrix $R_\varepsilon(\boldsymbol{\theta})$ satisfies*

$$\log|R_\varepsilon(\boldsymbol{\theta})| = \log|D_\varepsilon(\boldsymbol{\theta})| + \log\left|I_n + W_S^\top D_\varepsilon(\boldsymbol{\theta})^{-1} W_S\right|.$$

*Equivalently,*

$$\log|R_\varepsilon(\boldsymbol{\theta})| = \sum_{j=1}^{d}\log(1-\theta_j^2+\varepsilon) + \log\left|I_n + W_S^\top D_\varepsilon(\boldsymbol{\theta})^{-1} W_S\right|.$$

*Moreover, when $n \ll d$, this decomposition reduces the computational cost of evaluating the log-determinant from the naive order $\mathcal{O}(d^3)$ to*

$$\mathcal{O}(n^2 d + n^3),$$

*which is dominated by $\mathcal{O}(n^2 d)$ whenever $n^3$ is negligible relative to $n^2 d$.*

*Proof* Since

$$D_\varepsilon(\boldsymbol{\theta}) = \operatorname{diag}(1-\theta_1^2+\varepsilon, \ldots, 1-\theta_d^2+\varepsilon),$$

and, under the assumptions of Theorem 1, its diagonal entries are strictly positive, $D_\varepsilon(\boldsymbol{\theta})$ is nonsingular. Applying the matrix determinant lemma to

$$R_\varepsilon(\boldsymbol{\theta}) = D_\varepsilon(\boldsymbol{\theta}) + W_S W_S^\top$$

yields

$$\begin{aligned}|R_\varepsilon(\boldsymbol{\theta})| &= \left|D_\varepsilon(\boldsymbol{\theta}) + W_S W_S^\top\right| \\ &= |D_\varepsilon(\boldsymbol{\theta})| \left|I_n + W_S^\top D_\varepsilon(\boldsymbol{\theta})^{-1} W_S\right|.\end{aligned}$$

Taking logarithms gives

$$\log|R_\varepsilon(\boldsymbol{\theta})| = \log|D_\varepsilon(\boldsymbol{\theta})| + \log\left|I_n + W_S^\top D_\varepsilon(\boldsymbol{\theta})^{-1} W_S\right|.$$

Because $D_\varepsilon(\boldsymbol{\theta})$ is diagonal, we also have

$$\log|D_\varepsilon(\boldsymbol{\theta})| = \sum_{j=1}^{d}\log(1-\theta_j^2+\varepsilon),$$

which proves the stated decomposition.

Next, we discuss the computational cost. A direct evaluation of

$$\log\left|D_\varepsilon(\boldsymbol{\theta}) + W_S W_S^\top\right|$$

requires computing the log-determinant of a $d \times d$ matrix.

Using, for example, a Cholesky factorization, this step has the naive computational cost $\mathcal{O}(d^3)$, in addition to the cost of forming the matrix itself. The latter consists of $\mathcal{O}(d)$ operations for computing the diagonal entries of $D_\varepsilon(\boldsymbol{\theta})$ and $\mathcal{O}(d^2 n)$ operations for forming $W_S W_S^\top$. Thus, a direct evaluation incurs a total cost of order

$$\mathcal{O}(d + d^2 n + d^3),$$

which, in high-dimensional settings where $n < d$, is typically dominated by $\mathcal{O}(d^3)$.

By contrast, the decomposition above only requires the determinant of the $n \times n$ matrix

$$I_n + W_S^\top D_\varepsilon(\boldsymbol{\theta})^{-1} W_S.$$

Forming

$$W_S^\top D_\varepsilon(\boldsymbol{\theta})^{-1} W_S$$

is equivalently to forming

$$Z_S^\top \tilde{D}_\varepsilon(\boldsymbol{\theta}) Z_S,$$

where $\tilde{D}_\varepsilon(\boldsymbol{\theta}) = \mathrm{diag}(\frac{\theta_1^2}{1-\theta_1^2+\varepsilon}, \dots, \frac{\theta_d^2}{1-\theta_d^2+\varepsilon})$. Since $\tilde{D}_\varepsilon(\boldsymbol{\theta})$ is diagonal, forming $\tilde{D}_\varepsilon(\boldsymbol{\theta}) Z_S$ costs $\mathcal{O}(dn)$ operations only, thereby forming $Z_S^\top \tilde{D}_\varepsilon(\boldsymbol{\theta}) Z_S$ costs $\mathcal{O}(n^2 d)$ operations, subsequently computing the summation of $I_n$ and the resulting $n \times n$ matrix, $Z_S^\top \tilde{D}_\varepsilon(\boldsymbol{\theta}) Z_S$, requires $\mathcal{O}(n)$ operations only as $I_n$ is diagonal.

In addition, computing the log-determinant of the resulting $n \times n$ matrix, $I_n + Z_S^\top \tilde{D}_\varepsilon(\boldsymbol{\theta}) Z_S$, costs $\mathcal{O}(n^3)$ operations via Cholesky factorization and the diagonal term

$$\sum_{j=1}^{d} \log(1 - \theta_j^2 + \varepsilon)$$

costs only $\mathcal{O}(d)$ operations.

Therefore, the total cost of the decomposed evaluation is

$$\mathcal{O}(d + dn + n^2 d + n + n^3) = \mathcal{O}(n^2 d + n^3).$$

When $n \ll d$, this is substantially smaller than the naive $\mathcal{O}(d^3)$ cost. In particular, if $n^3 = o(n^2 d)$, then the leading order is $\mathcal{O}(n^2 d)$. □

**Lemma 3** (Trace decomposition) *Under the notation and assumptions of Theorem 1, the trace term satisfies*

$$\begin{aligned}\mathrm{tr}\Big(S\Sigma(\boldsymbol{\theta}, \boldsymbol{\sigma}^2)^{-1}\Big) &= \mathrm{tr}\Big(\widetilde{S}_\sigma\, R_\varepsilon(\boldsymbol{\theta})^{-1}\Big)\\ &= \frac{1}{n}\,\mathrm{tr}\Big(\widetilde{Z}_\sigma\, R_\varepsilon(\boldsymbol{\theta})^{-1} \widetilde{Z}_\sigma^\top\Big),\end{aligned}$$

*where* $\widetilde{S}_\sigma = D_\sigma^{-1/2}\, S\, D_\sigma^{-1/2}$, $\widetilde{Z}_\sigma = Y D_\sigma^{-1/2}$. *Moreover, when* $n \ll d$, *this decomposition reduces the computational cost of evaluating the trace term from the naive order* $\mathcal{O}(d^3)$ *to*

$$\mathcal{O}(n^2 d + n^3),$$

*which is dominated by* $\mathcal{O}(n^2 d)$ *whenever* $n^3$ *is negligible relative to* $n^2 d$.

*Proof* According to the commutativity of trace operation with the decomposition of $\Sigma(\boldsymbol{\theta}, \boldsymbol{\sigma}^2)$, we have

$$\operatorname{tr}\Big(S\Sigma(\boldsymbol{\theta}, \boldsymbol{\sigma}^2)^{-1}\Big) = \operatorname{tr}\Big(SD_\sigma^{-1/2} R_\varepsilon(\boldsymbol{\theta})^{-1} D_\sigma^{-1/2}\Big) = \operatorname{tr}\Big(\widetilde{S}_\sigma\, R_\varepsilon(\boldsymbol{\theta})^{-1}\Big).$$

where $\widetilde{S}_\sigma = D_\sigma^{-1/2}\, S\, D_\sigma^{-1/2}$.

Under the notation and assumptions of Theorem 1, the matrix $\widetilde{S}_\sigma$ satisfies

$$\widetilde{S}_\sigma = \frac{1}{n} \widetilde{Z}_\sigma^\top \widetilde{Z}_\sigma,$$

where $\widetilde{Z}_\sigma = Y D_\sigma^{-1/2}$. Thus we obtain the compressed form:

$$\operatorname{tr}\Big(\widetilde{S}_\sigma\, R_\varepsilon(\boldsymbol{\theta})^{-1}\Big) = \frac{1}{n} \operatorname{tr}\Big(\widetilde{Z}_\sigma\, R_\varepsilon(\boldsymbol{\theta})^{-1} \widetilde{Z}_\sigma^\top\Big).$$

which proves the stated formula.

We next analyze the computational cost. Computing the inverse of $\Sigma(\boldsymbol{\theta}, \boldsymbol{\sigma}^2)^{-1}$ has the naive computational cost $\mathcal{O}(d^3)$, while forming a matrix as $S\Sigma(\boldsymbol{\theta}, \boldsymbol{\sigma}^2)^{-1}$ costs $\mathcal{O}(d^3)$. In addition, the trace computing is an $\mathcal{O}(d^2)$ operation. Thus, a direct evaluation incurs a total cost of order

$$\mathcal{O}(d^3 + d^2)$$

which is dominated by $\mathcal{O}(d^3)$. By contrast, similar to Lemma 1, we apply the Woodbury identity to

$$R_\varepsilon(\boldsymbol{\theta}) = D_\varepsilon(\boldsymbol{\theta}) + W_S W_S^\top,$$

which yields an inverse expressed through the diagonal matrix $D_\varepsilon(\boldsymbol{\theta})^{-1}$ and an $n \times n$ matrix

$$I_n + W_S^\top D_\varepsilon(\boldsymbol{\theta})^{-1} W_S.$$

where $D_\varepsilon(\boldsymbol{\theta})^{-1}$ costs $\mathcal{O}(d)$ up to forming diagonal entries, and forming $W_S^\top D_\varepsilon(\boldsymbol{\theta})^{-1} W_S$ costs $\mathcal{O}(n^2 d)$. In addition, computing the $n \times n$ inverse costs $\mathcal{O}(n^3)$ followed by the $\mathcal{O}(n^2)$ trace operation. Hence the total cost of the trace term is

$$\mathcal{O}(d + n^2 + n^2 d + n^3) = \mathcal{O}(n^2 d + n^3),$$

When $n \ll d$, this is substantially smaller than the naive $\mathcal{O}(d^3)$ cost. In particular, if $n^3 = o(n^2 d)$, then the leading order is $\mathcal{O}(n^2 d)$. □

**Theorem 4** (Conditioning bounds via Woodbury) *Under the notation and assumptions of Theorem 1, let $u_j = 1 - \theta_j^2$ for $j = 1, \ldots, d$. For an arbitrary real square matrix $A$, let $\lambda_{\min}(A)$ and $\lambda_{\max}(A)$ denote its smallest and largest eigenvalues, respectively.*

*Then*

$$\lambda_{\min}\{R_\varepsilon(\boldsymbol{\theta})\} \geq \min_{1 \leq j \leq d} (u_j + \varepsilon),$$

*and*

$$\lambda_{\max}\{R_\varepsilon(\boldsymbol{\theta})\} \leq \max_{1 \leq j \leq d} (u_j + \varepsilon) + \|W_S\|_2^2.$$

*Consequently, the condition number satisfies*

$$\kappa\{R_\varepsilon(\boldsymbol{\theta})\} \leq \frac{\max_{1 \leq j \leq d}(u_j + \varepsilon) + \|W_S\|_2^2}{\min_{1 \leq j \leq d}(u_j + \varepsilon)}.$$

*Proof* We establish the lower and upper eigenvalue bounds separately.

We first prove the lower bound. For any $\boldsymbol{\nu} \in \mathbb{R}^d$, we have

$$\begin{aligned}\boldsymbol{\nu}^\top R_\varepsilon(\boldsymbol{\theta})\boldsymbol{\nu} &= \boldsymbol{\nu}^\top D_\varepsilon(\boldsymbol{\theta})\boldsymbol{\nu} + \boldsymbol{\nu}^\top W_S W_S^\top \boldsymbol{\nu} \\ &= \sum_{j=1}^{d} (u_j + \varepsilon)\nu_j^2 + \|W_S^\top \boldsymbol{\nu}\|_2^2.\end{aligned}$$

Since $\|W_S^\top \boldsymbol{\nu}\|_2^2 \geq 0$, it follows that

$$\boldsymbol{\nu}^\top R_\varepsilon(\boldsymbol{\theta})\boldsymbol{\nu} \geq \sum_{j=1}^{d} (u_j + \varepsilon)\nu_j^2 \geq \min_{1\leq j\leq d} (u_j + \varepsilon)\|\boldsymbol{\nu}\|_2^2.$$

Moreover, $R_\varepsilon(\boldsymbol{\theta})$ is real symmetric. Therefore, by the Rayleigh–Ritz characterization,

$$\lambda_{\min}\{R_\varepsilon(\boldsymbol{\theta})\} = \min_{\boldsymbol{\nu}\neq \mathbf{0}} \frac{\boldsymbol{\nu}^\top R_\varepsilon(\boldsymbol{\theta})\boldsymbol{\nu}}{\|\boldsymbol{\nu}\|_2^2} \geq \min_{1\leq j\leq d} (u_j + \varepsilon).$$

We next prove the upper bound. By Weyl's inequality,

$$\lambda_{\max}\{D_\varepsilon(\boldsymbol{\theta}) + W_S W_S^\top\} \leq \lambda_{\max}\{D_\varepsilon(\boldsymbol{\theta})\} + \lambda_{\max}(W_S W_S^\top).$$

Since $D_\varepsilon(\boldsymbol{\theta})$ is diagonal,

$$\lambda_{\max}\{D_\varepsilon(\boldsymbol{\theta})\} = \max_{1\leq j\leq d} (u_j + \varepsilon).$$

Furthermore, using the standard relationship between the spectral norm and the largest eigenvalue,

$$\lambda_{\max}\{W_S W_S^\top\} = \|W_S\|_2^2.$$

Consequently,

$$\lambda_{\max}\{R_\varepsilon(\boldsymbol{\theta})\} \leq \max_{1\leq j\leq d} (u_j + \varepsilon) + \|W_S\|_2^2.$$

Combining the lower and upper bounds gives

$$\kappa\{R_\varepsilon(\boldsymbol{\theta})\} = \frac{\lambda_{\max}\{R_\varepsilon(\boldsymbol{\theta})\}}{\lambda_{\min}\{R_\varepsilon(\boldsymbol{\theta})\}} \leq \frac{\max_{1\leq j\leq d}(u_j + \varepsilon) + \|W_S\|_2^2}{\min_{1\leq j\leq d}(u_j + \varepsilon)}.$$

This completes the proof. □

**Theorem 5** (Posterior propriety) *Under the notation and assumptions of Theorem 1, suppose that the probability model is specified as in Eq. 1 (in Section Method). As described in Section Method, assume that the calibration parameters $\theta_j$ are determined by the global shrinkage parameter $\tau$ and the local shrinkage parameters $\lambda_j$, $j = 1, \ldots, d$, with the latter assigned proper half-Cauchy priors. Further assume that the variance components satisfy*

$$\sigma_j^2 \sim \mathrm{IG}(a, b), \qquad a > 0, \quad b > 0, \quad j = 1, \ldots, d.$$

*Then, for any fixed $\varepsilon > 0$, the marginal normalizing constant*

$$m(Y) = \int p(Y \mid \tau, \boldsymbol{\lambda}, \boldsymbol{\sigma}^2)\pi(\tau) \prod_{j=1}^{d} \pi(\lambda_j) \prod_{j=1}^{d} \pi(\sigma_j^2)\, d\boldsymbol{\sigma}^2\, d\tau\, d\boldsymbol{\lambda}$$

*is finite. Consequently, the posterior distribution is proper.*

*Proof* For any $\varepsilon > 0$, Theorem 4 implies that

$$\lambda_{\min}\{R_\varepsilon(\boldsymbol{\theta})\} \geq \min_{1\leq j\leq d}(\mu_j + \varepsilon).$$

As stated in Theorem 4, we have $\mu_j = 1 - \theta_j \geq 0$ for all $j = 1, \ldots, d$. Therefore,

$$\mu_j + \varepsilon \geq \varepsilon, \qquad j = 1, \ldots, d,$$

and hence

$$\lambda_{\min}\{R_\varepsilon(\boldsymbol{\theta})\} \geq \varepsilon.$$

Consequently

$$|R_\varepsilon(\boldsymbol{\theta})| \geq \varepsilon^d,$$

and therefore,

$$|R_\varepsilon(\boldsymbol{\theta})|^{-n/2} \leq \varepsilon^{-nd/2}. \tag{1}$$

According to the probability model and the proposed parametrized structure for $\Sigma(\boldsymbol{\theta})$, the likelihood can be written as

$$p(Y \mid \boldsymbol{\theta}, \boldsymbol{\sigma}^2) = (2\pi)^{-nd/2}|D_\sigma|^{-n/2}|R_\varepsilon(\boldsymbol{\theta})|^{-n/2} \times \exp\left\{-\frac{n}{2}\operatorname{tr}\left(SD_\sigma^{-1/2}R_\varepsilon(\boldsymbol{\theta})^{-1}D_\sigma^{-1/2}\right)\right\},$$

where

$$D_\sigma = \operatorname{diag}(\sigma_1^2, \ldots, \sigma_d^2).$$

Next, we bound the exponential term. Define

$$A = \left[\Sigma_\varepsilon(\boldsymbol{\theta}, \boldsymbol{\sigma}^2)\right]^{-1} = D_\sigma^{-1/2}R_\varepsilon(\boldsymbol{\theta})^{-1}D_\sigma^{-1/2}.$$

By Theorem 1, $\Sigma_\varepsilon(\boldsymbol{\theta}, \boldsymbol{\sigma}^2)$ is positive definite. Therefore, by standard properties of positive definite matrices, $A$ is positive definite, and so is its principal square root $A^{1/2}$.

Moreover, since $S$ is positive semidefinite and positive semidefiniteness is preserved under congruence transformations by nonsingular matrices, it follows that

$$A^{1/2}SA^{1/2}$$

is positive semidefinite. Moreover, by the cyclic invariance of the trace,

$$\operatorname{tr}(SA) = \operatorname{tr}(SA^{1/2}A^{1/2}) = \operatorname{tr}(A^{1/2}SA^{1/2}).$$

Since the trace of a positive semidefinite matrix is nonnegative, we obtain

$$\operatorname{tr}(SA) = \operatorname{tr}(A^{1/2}SA^{1/2}) \geq 0.$$

Therefore,

$$\exp\left\{-\frac{n}{2}\operatorname{tr}(SA)\right\} \leq 1 \tag{2}$$

Combining the upper bound for the exponential term in Eq. (2) with the determinant bound in Eq. (1), we obtain

$$p(Y \mid \boldsymbol{\theta}, \boldsymbol{\sigma}^2) \leq (2\pi)^{-nd/2}\varepsilon^{-nd/2}|D_\sigma|^{-n/2}.$$

Let

$$C_\varepsilon = (2\pi)^{-nd/2}\varepsilon^{-nd/2}.$$

For any fixed $\varepsilon > 0$, we have $C_\varepsilon < \infty$.

Since

$$|D_\sigma|^{-n/2} = \prod_{j=1}^{d}(\sigma_j^2)^{-n/2},$$

it follows that

$$p(Y \mid \boldsymbol{\theta}, \boldsymbol{\sigma}^2) \le C_\varepsilon \prod_{j=1}^{d} (\sigma_j^2)^{-n/2}. \tag{3}$$

Substituting Eq. (3) into the marginal normalizing constant yields

$$m(Y) \le C_\varepsilon \left( \int_0^\infty \pi(\tau)\, d\tau \right) \prod_{j=1}^{d} \left[ \int_0^\infty \pi(\lambda_j)\, d\lambda_j \right]$$
$$\times \prod_{j=1}^{d} \left[ \int_0^\infty (\sigma_j^2)^{-n/2} \pi(\sigma_j^2)\, d\sigma_j^2 \right].$$

The half-Cauchy priors assigned to $\tau$ and $\lambda_j$ are proper. Therefore,

$$\int_0^\infty \pi(\tau)\, d\tau < \infty, \qquad \int_0^\infty \pi(\lambda_j)\, d\lambda_j < \infty, \qquad j = 1, \ldots, d.$$

It remains to verify that each variance-related integral is finite. Since

$$\sigma_j^2 \sim \mathrm{IG}(a, b), \qquad a > 0, \quad b > 0,$$

the density of $\sigma_j^2$ is

$$\pi(\sigma_j^2) = \frac{b^a}{\Gamma(a)} (\sigma_j^2)^{-a-1} \exp\left( -\frac{b}{\sigma_j^2} \right), \qquad \sigma_j^2 > 0.$$

Hence,

$$\int_0^\infty (\sigma_j^2)^{-n/2} \pi(\sigma_j^2)\, d\sigma_j^2 = \frac{b^a}{\Gamma(a)} \int_0^\infty (\sigma_j^2)^{-a-n/2-1} \exp\left( -\frac{b}{\sigma_j^2} \right) d\sigma_j^2.$$

Using the change of variables

$$x = \frac{b}{\sigma_j^2}, \qquad \sigma_j^2 = \frac{b}{x}, \qquad d\sigma_j^2 = -b x^{-2}\, dx,$$

we obtain

$$\begin{aligned} &\int_0^\infty (\sigma_j^2)^{-a-n/2-1} \exp\left( -\frac{b}{\sigma_j^2} \right) d\sigma_j^2 \\ &= b^{-a-n/2} \int_0^\infty x^{a+n/2-1} e^{-x}\, dx \\ &= b^{-a-n/2} \Gamma(a + n/2). \end{aligned}$$

Consequently,

$$\int_0^\infty (\sigma_j^2)^{-n/2} \pi(\sigma_j^2)\, d\sigma_j^2 = \frac{b^{-n/2} \Gamma(a + n/2)}{\Gamma(a)} < \infty.$$

Since all factors in the upper bound for $m(Y)$ are finite, we conclude that

$$m(Y) < \infty.$$

Therefore, the posterior distribution is proper. □

**Theorem 6** (Oracle approximation bound under active-node sparsity) *Under the notation and assumptions of Theorem 1, let $R^*$ denote the population correlation matrix and let $R_S$ be its empirical counterpart (i.e., a borrowed pilot correlation matrix). Suppose that, on an event $\mathcal{E}_n$, the sample correlation matrix satisfies*

$$\|R_S - R^*\|_F \le C_0\sqrt{\frac{m^* \log d}{n}},$$

*where $m^*$ denotes the number of nonzero off-diagonal entries in $R^*$.*

*Define the oracle approximation error*

$$a^*_{n,\varepsilon} = \inf_{\boldsymbol{\theta}\in\Theta} \|R_\varepsilon(\boldsymbol{\theta}) - R^*\|_F.$$

*Assume that $\widehat{\boldsymbol{\theta}}$ satisfies the approximate projection condition*

$$\|R_\varepsilon(\widehat{\boldsymbol{\theta}}) - R_S\|_F \le \inf_{\boldsymbol{\theta}\in\Theta} \|R_\varepsilon(\boldsymbol{\theta}) - R_S\|_F + r_n,$$

*where $r_n \ge 0$ is the optimization or approximation tolerance. Then, on $\mathcal{E}_n$,*

$$\|R_\varepsilon(\widehat{\boldsymbol{\theta}}) - R^*\|_F \le a^*_{n,\varepsilon} + 2C_0\sqrt{\frac{m^* \log d}{n}} + r_n.$$

*In particular, if the support of $R^*$ is induced by at most $s$ active nodes, then*

$$m^* \le \frac{s(s-1)}{2},$$

*and hence*

$$\|R_\varepsilon(\widehat{\boldsymbol{\theta}}) - R^*\|_F \le a^*_{n,\varepsilon} + 2C_0\sqrt{\frac{s(s-1)\log d}{2n}} + r_n.$$

*Proof* We decompose the estimation error into an approximation component, a stochastic component, and an optimization tolerance component.

By the triangle inequality,

$$\|R_\varepsilon(\widehat{\boldsymbol{\theta}}) - R^*\|_F \le \|R_\varepsilon(\widehat{\boldsymbol{\theta}}) - R_S\|_F + \|R_S - R^*\|_F.$$

Using the approximate projection condition, we obtain

$$\|R_\varepsilon(\widehat{\boldsymbol{\theta}}) - R^*\|_F \le \inf_{\boldsymbol{\theta}\in\Theta} \|R_\varepsilon(\boldsymbol{\theta}) - R_S\|_F + r_n + \|R_S - R^*\|_F.$$

It remains to control the empirical projection term. For any $\boldsymbol{\theta} \in \Theta$, another application of the triangle inequality gives

$$\|R_\varepsilon(\boldsymbol{\theta}) - R_S\|_F \le \|R_\varepsilon(\boldsymbol{\theta}) - R^*\|_F + \|R_S - R^*\|_F.$$

Taking the infimum over $\boldsymbol{\theta} \in \Theta$ on both sides yields

$$\begin{aligned}\inf_{\boldsymbol{\theta}\in\Theta} \|R_\varepsilon(\boldsymbol{\theta}) - R_S\|_F &\le \inf_{\boldsymbol{\theta}\in\Theta} \|R_\varepsilon(\boldsymbol{\theta}) - R^*\|_F + \|R_S - R^*\|_F \\ &= a^*_{n,\varepsilon} + \|R_S - R^*\|_F.\end{aligned}$$

Combining the preceding inequalities gives

$$\|R_\varepsilon(\widehat{\boldsymbol{\theta}}) - R^*\|_F \le a^*_{n,\varepsilon} + 2\|R_S - R^*\|_F + r_n.$$

On the event $\mathcal{E}_n$, by assumption,

$$\|R_S - R^*\|_F \le C_0\sqrt{\frac{m^* \log d}{n}}.$$

Therefore,

$$\|R_\varepsilon(\widehat{\boldsymbol{\theta}}) - R^*\|_F \le a^*_{n,\varepsilon} + 2C_0\sqrt{\frac{m^* \log d}{n}} + r_n.$$

Finally, if the support of $R^*$ is induced by at most $s$ active nodes, then the maximum number of possible nonzero off-diagonal entries among these active nodes is

$$\binom{s}{2} = \frac{s(s-1)}{2}.$$

Thus,

$$m^* \le \frac{s(s-1)}{2}.$$

Substituting this bound into the preceding inequality gives

$$\|R_\varepsilon(\widehat{\boldsymbol{\theta}}) - R^*\|_F \le a^*_{n,\varepsilon} + 2C_0\sqrt{\frac{s(s-1)\log d}{2n}} + r_n.$$

This completes the proof. □

**Theorem 7** *Under the notation and assumptions of Theorem 6, define*

$$a_n := C_0\sqrt{\frac{m^* \log d}{n}},$$

*where $C_0 > 0$ is a constant. Let $R_\varphi$ be a reconstructed correlation matrix associated with $R_S$ (enlarged or shrunk) satisfying*

$$R_\varphi[ij] = \varphi\, R_S[ij], \quad i \ne j, \qquad R_\varphi[jj] = 1, \quad j = 1, \dots, d.$$

*Assume that the parameter $\varphi > 0$ satisfies*

$$\varphi = 1 + o\left(\frac{1}{d}\right), \qquad |\varphi\, R_S[ij]| \le 1, \quad i \ne j.$$

*Suppose that*

$$|R_S - R^*\|_F \le a_n,$$

*and that the construction of $R_\varphi$ yields the perturbation bound*

$$|R_\varphi - R^*\|_F \le \varphi a_n + |1-\varphi|\sqrt{d^2 - d}.$$

*Then, for all sufficiently large $d$, there exists a constant $K > 0$ such that*

$$|R_\varphi - R^*\|_F \le \left(1 + \frac{K}{d}\right) a_n + K\sqrt{1 - \frac{1}{d}}.$$

*More precisely, let*

$$\varphi = 1 + \varepsilon_d, \qquad d|\varepsilon_d| \longrightarrow 0.$$

*Then*

$$|R_\varphi - R^*\|_F \le (1 + |\varepsilon_d|)a_n + \big(d|\varepsilon_d|\big)\sqrt{1 - \frac{1}{d}}.$$

*Consequently,*

$$|R_\varphi - R^*\|_F \le a_n + o(1) + o\left(\sqrt{\frac{m^* \log d}{n}}\right).$$

*In particular, if*

$$\frac{m^* \log d}{n} \longrightarrow 0,$$

*then*

$$|R_\varphi - R^*\|_F \longrightarrow 0.$$

*Proof* Let

$$a_n = C_0\sqrt{\frac{m^* \log d}{n}}.$$

By the assumed perturbation bound,

$$|R_\varphi - R^*\|_F \le \varphi a_n + |1-\varphi|\sqrt{d^2-d}.$$

Substituting the definition of $a_n$ gives

$$\|R_\varphi - R^*\|_F \le \varphi C_0\sqrt{\frac{m^* \log d}{n}} + |1-\varphi|\sqrt{d^2-d}. \tag{4}$$

Since $\varphi = 1 + o(1/d)$, there exists a sequence $\{\varepsilon_d\}$ such that

$$\varphi = 1 + \varepsilon_d, \qquad d\varepsilon_d \longrightarrow 0.$$

Thus $|1-\varphi| = |\varepsilon_d|$, and Eq. 4 becomes

$$|R_\varphi - R^*\|_F \le (1+\varepsilon_d)C_0\sqrt{\frac{m^* \log d}{n}} + |\varepsilon_d|\sqrt{d^2-d}. \tag{5}$$

Noting that $\sqrt{d^2-d} = d\sqrt{1-1/d}$, we have

$$|\varepsilon_d|\sqrt{d^2-d} = \big(d|\varepsilon_d|\big)\sqrt{1-\frac{1}{d}}.$$

Using $1+\varepsilon_d \le 1 + |\varepsilon_d|$, it follows from Eq. 5 that

$$|R_\varphi - R^*\|_F \le \big(1+|\varepsilon_d|\big)C_0\sqrt{\frac{m^* \log d}{n}} + \big(d|\varepsilon_d|\big)\sqrt{1-\frac{1}{d}}. \tag{6}$$

Because $d|\varepsilon_d| \to 0$, the sequence $\{d|\varepsilon_d|\}$ is bounded. Hence there exists a constant $K > 0$ such that, for all sufficiently large $d$,

$$d|\varepsilon_d| \le K, \qquad |\varepsilon_d| \le \frac{K}{d}.$$

Substituting these into Eq. 6 yields

$$\|R_\varphi - R^*\|_F \le \Big(1+\frac{K}{d}\Big)C_0\sqrt{\frac{m^* \log d}{n}} + K\sqrt{1-\frac{1}{d}},$$

which establishes the finite-dimensional bound.

To derive the sharper asymptotic bound, expand Eq. 6 as

$$\|R_\varphi - R^*\|_F \le C_0\sqrt{\frac{m^* \log d}{n}} + |\varepsilon_d|C_0\sqrt{\frac{m^* \log d}{n}} + \big(d|\varepsilon_d|\big)\sqrt{1-\frac{1}{d}}.$$

Since $|\varepsilon_d| = o(1/d)$ and $d|\varepsilon_d| = o(1)$, we obtain

$$|\varepsilon_d|C_0\sqrt{\frac{m^* \log d}{n}} = o\left(\sqrt{\frac{m^* \log d}{n}}\right), \qquad \big(d|\varepsilon_d|\big)\sqrt{1-\frac{1}{d}} = o(1).$$

Therefore,

$$\|R_\varphi - R^*\|_F \le C_0\sqrt{\frac{m^* \log d}{n}} + o\left(\sqrt{\frac{m^* \log d}{n}}\right) + o(1).$$

Finally, if $(m^* \log d)/n \to 0$, then $C_0\sqrt{m^* \log d/n} \to 0$, and the preceding bound gives

$$\|R_\varphi - R^*\|_F \longrightarrow 0.$$

□

*Remark 1* The Frobenius-norm error bound consists of two components:

$$C_0\sqrt{\frac{m^* \log d}{n}} \quad \text{and} \quad |1-\varphi|\sqrt{d^2-d}.$$

The first term represents the estimation error associated with the sample correlation matrix $R_S$, where $n$ denotes the sample size. The second term quantifies the perturbation induced by the rescaling parameter $\varphi$ and depends on the matrix dimension $d$.

Since $\sqrt{d^2-d} = d\sqrt{1-1/d}$, the perturbation term can be written as

$$|1-\varphi|\sqrt{d^2-d} = d|\varphi-1|\sqrt{1-\frac{1}{d}} < d|\varphi-1|.$$

Thus, the condition $\varphi = 1 + o(1/d)$ is sufficient to guarantee that $d|\varphi-1| \to 0$, so that the cumulative rescaling perturbation vanishes asymptotically. The parameter $\varphi$ need not be greater than 1 or equal to 1; it may approach 1 from either side.

If this condition is not satisfied but the weaker condition $\varphi = 1 + O\left(\frac{1}{d}\right)$ holds, then $d|\varphi-1| = O(1)$. In this case, the cumulative perturbation remains bounded.

### Supplementary Note 2. The details of the covariance structures and the four MSE measures

This section covers the details of the covariance structures and the four MSE measures used in simulation studies.

#### Supplementary Note 2.1. Details of covariance structures

Here, we provide details of the three covariance structures—block, banded, and Toeplitz—used in the simulation studies, following [1–3].

1. **Block.** The indices $1, 2, \ldots, d$ were partitioned into five ordered groups of equal size. Correlations were set to be moderate within each group and weak across groups, thereby mimicking clustered dependence patterns frequently encountered in brain connectivity studies:

$$\Sigma^*_{ij} = \begin{cases} 1, & i = j, \\ 0.6, & i \text{ and } j \text{ belong to the same group}, \\ 0, & \text{otherwise}. \end{cases}$$

2. **Banded.** Two variables indexed by $i$ and $j$ were conditionally connected when $|i-j| \le 3$, yielding the covariance structure

$$\Sigma^*_{ij} = \begin{cases} 1 - \dfrac{|i-j|}{3}, & |i-j| \le 3, \\ 0, & \text{otherwise}. \end{cases}$$

3. **Toeplitz.** Under the Toeplitz structure, covariance entries depended only on $|i-j|$, so that all elements along the same diagonal were equal:

$$\Sigma^*_{ij} = 0.75^{|i-j|}.$$

#### Supplementary Note 2.2. Four MSE measures

Four MSE measures were also used in the simulation studies to assess the effectiveness of the shrinkage terms and were calculated as belows, following [4]:

$$\mathrm{MSE}_{\mathcal{P}}^{S} = \frac{1}{|\mathcal{P}|} \sum_{(i,j)\in\mathcal{P}} (R_{ij}^{S} - R_{ij}^{*})^2 \tag{7}$$

$$\mathrm{MSE}_{\mathcal{Q}}^{S} = \frac{1}{|\mathcal{Q}|} \sum_{(i,j)\in\mathcal{Q}} (R_{ij}^{S} - R_{ij}^{*})^2 \tag{8}$$

and

$$\mathrm{MSE}_{\mathcal{P}}^{\mathrm{IBB}} = \frac{1}{|\mathcal{P}|} \sum_{(i,j)\in\mathcal{P}} (\hat{R}_{ij} - R_{ij}^{*})^2 \tag{9}$$

$$\mathrm{MSE}_{\mathcal{Q}}^{\mathrm{IBB}} = \frac{1}{|\mathcal{Q}|} \sum_{(i,j)\in\mathcal{Q}} (\hat{R}_{ij} - R_{ij}^{*})^2 \tag{10}$$

where $R_{ij}^{S}$, $\hat{R}_{ij}$ and $R_{ij}^{*}$ denote the entry of the sample correlation, posterior mean correlation and underlying true correlation matrix, respectively and

$$\mathcal{P} = \{(i,j) : |\hat{R}_{ij} - R_{ij}^{S}| > \delta\},$$

denotes the index set of entries whose absolute differences exceed a given threshold $\delta$, and $\mathcal{Q}$ denotes the complement of $\mathcal{P}$. We set $\delta = 0.05$ to represent a practically meaningful level of correction [5].

#### Supplementary Note 3. Gaussian-likelihood-based classification

To assess downstream utility in the semi-synthetic study, we further evaluated a Gaussian-likelihood-based classification task. For each repetition, we performed 3-fold cross-validation on the semi-synthetic dataset. In each split, we used the training folds (with known group labels) to fit two Gaussian models, one for each group, and estimated the corresponding covariance matrices. Because our primary interest was covariance estimation, we set the group mean vectors $\widehat{\mu}_G$ and $\widehat{\mu}_C$ to the sample means of the training folds to simplify computation. We set the class prior to the empirical class proportion in the training fold, $\widehat{\pi}_g = \frac{n_g}{n_{\mathrm{train}}}$. Similar to [6], for a validation observation $\mathbf{y}$, we computed an unnormalized log-posterior score for each group:

$$\log p(g \mid \mathbf{y}) = \log \mathcal{N}(\mathbf{y}; \widehat{\mu}_g, \widehat{\Sigma}_g) + \log \widehat{\pi}_g. \tag{11}$$

We then assigned $\mathbf{y}$ to the group with the larger score.

### Supplementary Note 4. Validation using the New York University Child Study Center dataset
